# Harnessing Immunoinformatic for HIV-2: Multi-Epitope Vaccine Design Targeting the env Protein (CAA28914.1)

**DOI:** 10.1101/2025.06.06.658249

**Authors:** Muhammad Ali, Yasrab Arafat, Sheikh Ahmed Gull, Shah Zainab, Fareeza Anwar, Qanbar Abbas

## Abstract

**Background:** HIV-2 is a major threat to human health because of its rising global incidence. As a result, the development of effective immunostimulatory molecules for vaccine design against HIV-2 has garnered increasing interest.

**Objective:** The main objective was to make a multi Targeting epitoptpic vaccine for HIV-2 envelope protein using immunoinformatic approaches.

**Methods:** Sequence were retrieved from NCBI. B & T-cell epitopes were predicted with assessments of antigenicity, allergenicity, and cytokine-inducing potential. Physicochemical properties, docking studies, and codon optimization for expression was also performed using different software tools.

**Findings:** The designed vaccine construct (477 amino acids, molecular weight of 54 kDa) showed a strong antigenicity with VaxiJen score of 0.7146 which was non-allergenic and non-toxic with favorable physicochemical parameters (Mw: ∼53 kDa; pI: 9.89; stable and soluble). The construct demonstrated broad population coverage for MHC-I (West Africa: 69.51%, Pakistan: 78%, Global: 82%) and MHC-II (Global: 99%). Out of 16 peptides analyzed, 7 were predicted as IL-10 inducers 2 as a strong IL-4 inducer, and 2 were IFN-γ inducing epitopes. Structural validation confirmed high-quality folding (96% residues in favored regions; ERRAT score: 84.35). Docking results revealed a strong binding affinity to TLR2 (PDB ID: 2AOZ; energy: -1481.1 kcal/mol). MD simulation confirmed structural stability (RMSD ∼1.37 nm; Rg ∼3.65 nm). Immune simulation predicted robust humoral and cellular responses with sustained antibody production and cytokine activation. Codon optimization is given to a Codon Adaptation Index (CAI) of 0.628 with a guanine (G) and cytosine (C) nucleotides content of 65.48%, suitable for *E. coli* expression.

**Conclusion:** This vaccine construct is immunogenic, stable, and able to elicit both antibody-mediated immunity and cell-mediated immunity responses. Furthermore, In-vitro or In-Vivo Validations are recommended to verify its effectiveness

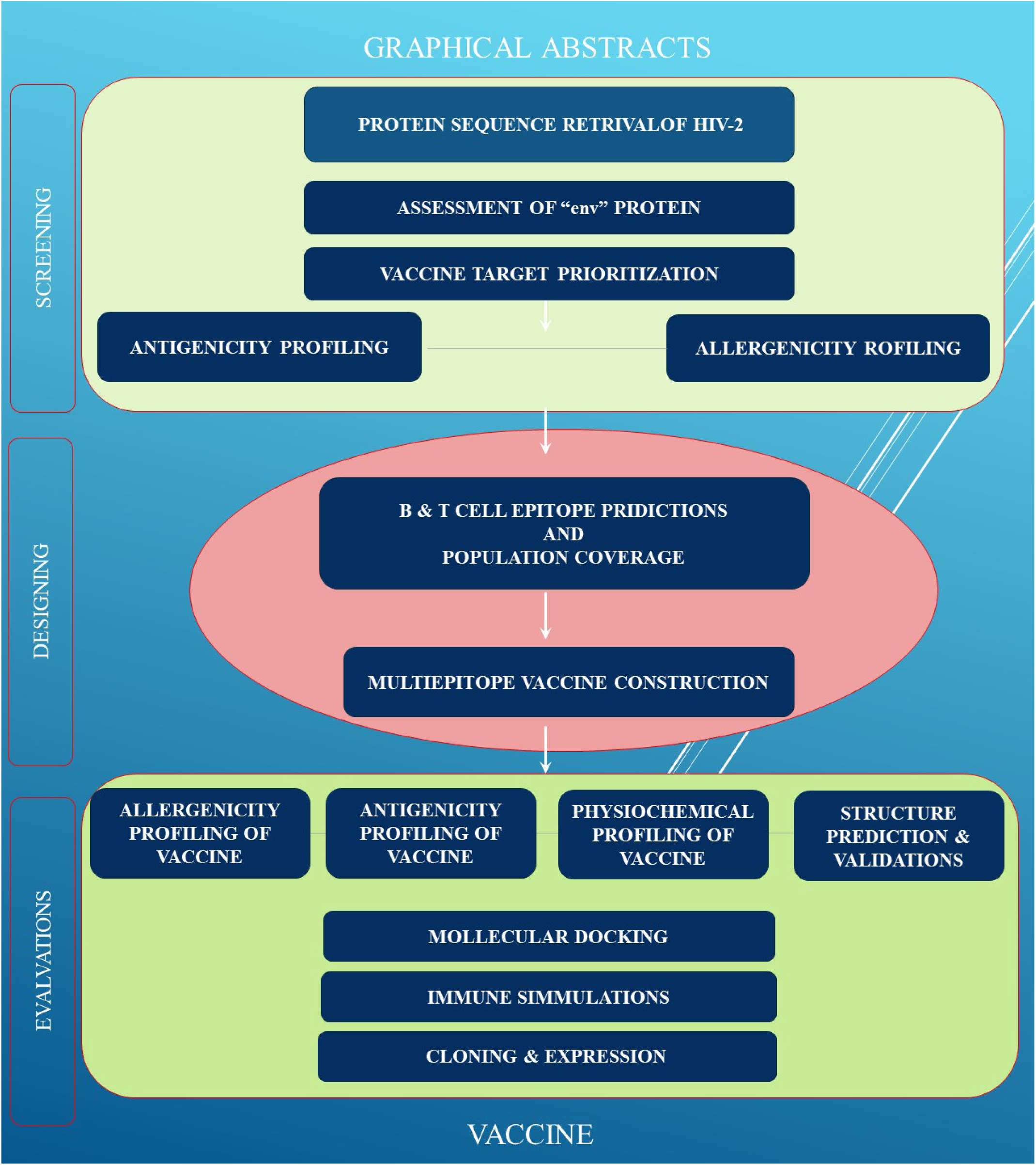

## Introduction

Human immunodeficiency virus type 2 (HIV-2) is primarily indigenous to West Africa but going up cases have been reported in differences counties like European continent, the Republic of India, and the United States of America due to globalization and migration. Despite its regional prevalence, HIV-2 remains underrecognized and often neglected in global HIV/AIDS control programs, which leads to unique challenges in diagnosis, patient care, and management (Gottlieb et al., 2018; Campbell et al., 2011). HIV-2 differs from HIV-1 in its natural history and disease progression. Evidence suggests that HIV-2 generally progresses more slowly than HIV-1, characterized by a longer asymptomatic phase and a slowing decrease in CD4 counts (Palm et al., 2022). Molecular studies expressed that intra-patient evolution of HIV-2 including changes in glycosylation sites and viral diversity can affect disease increases, with slower decreases showing unique molecular patterns compared to speeded up. HIV-2 gives different challenges for antiretroviral therapy due to drug vulnerability and resistance profiles compared to HIV-1. Optimal treatment has remained debatable till now, due to by no major clinical trials on HIV-2, which limits treatment guidelines. Correct diagnostics that can make HIV-2 different from HIV-1 are essential for proper management but remain difficult to do particularly in resource-constrained settings (Gottlieb et al., 2018; Campbell et al., 2011).

The HIV-2 env gene is essential for viral entry into host cells and central element in immune evasion. The Env protein is a major target for vaccine development; however, its structural complexity and immune escape mechanisms pose significant challenges for effective immunization strategies (Antanasijević et al., 2020; Zhao et al., 2022). Compared to HIV-1, HIV-2 infection is characterized by notably lower viral loads and less efficient transmission, resulting in a longer incubation period and slower increment to AIDS (Williams et al., 2023). Animal model studies have shown that HIV-2 replicates less than HIV-1, which contributes to its lower pathogenicity (Agarwal et al., 2020). Immuno-informatics use computational and bioinformatics approaches to analyze immune system components, host-pathogen interactions, and vaccine design processes. This field shed light on the identification of antigenic epitopes from genetic and protein sequence data that can elicit protective immune responses. By combining data from genomics, proteomics, and structural biology, immune informatics provides crucial insights that go up and optimize vaccine development (Oli et al., 2020; Joon et al., 2022). Epitope-based vaccines stimulate broad immune activation and good immunity by inducing robust antibody production via B cells and activating both cytotoxic and helper T cells. This dual role and activation contributes to on spot defense and long-term immune memory (Zhang, 2017; Waqas et al., 2022). Incorporating adjuvants and optimized linkers enhances immunogenicity and prolongs the immune response. Moreover, computational design facilitates the selection of epitopes hat are bound to a diverse array of human leukocyte antigen (HLA) alleles, ensuring broader population coverage and vaccine effectiveness across genetically heterogeneous groups (Zhang, 2017; Waqas et al., 2022).

To Targeting conserved regions within the HIV-2 env protein is a good strategy to overcome challenges given by viral mutation and antigenic variability. Multi-epitope vaccines can focus on these stable regions to induce broader and more durable immune protection (Zhang, 2017; Ahmad et al., 2024). Immuno-informatics use in-silico tools for the precise identification of immunodominant, conserved epitopes for vaccine (Yang et al., 2020). These computational methods provided the methods vaccine design by predicting high-affinity binding sites and optimizing antigen presentation, which is important for rapidly evolving viruses such as HIV-2. In our study, we design a multi-epitope HIV-2 Env protein vaccine candidate (CAA28914.1) using immune-informatics. We use computational tools to find conserved and immunogenic epitopes, integrate them into a subunit vaccine, and provide an in-silico process for experimental validation.

## Methods and Materials

For the Designing and building the multi-epitope vaccine candidate involved these following steps.

### Extraction of protein sequences

The reference sequence of the HIV-2 envelope (env) protein, with accession number >CAA28914.1, was retrieved from the National Center for Biotechnology Information (NCBI) database (available at: https[:]//www[.]ncbi[.]nlm[.]nih[.]gov). This sequence serves as a representative entry for the HIV-2 env gene product and was selected for further bioinformatic analysis. (Pruitt, Tatusova, & Maglott, 2005).

### Assessment of Selected env Protein

The physicochemical characteristics of the HIV-2 envelope glycoprotein (accession number CAA28914.1) were analyzed using the ProtParam tool available on the ExPASy server (accessible at: https[:]//web[.]expasy[.]org/protparam/). This computational tool gives these parameters; protein’s molecular weight, theoretical isoelectric point (pI), instability index, aliphatic index, and the Grand Average of Hydropathicity (GRAVY). These features explain the protein’s biochemical behavior, potential stability, and hydrophobic or hydrophilic nature, essential for understanding its structural and functional properties.

### B and T Cell Linear Epitope Forecasting

For the prediction of B cell and T cell epitopes of the HIV-2 env protein we used IEDB web server (accessible at: https[:]//www[.]iedb[.]org) with the use of the Bepipred Linear-Epitope Prediction tool (versions 1.0 & 2.0) (available at: https[:]//services[.]healthtech[.]dtu[.]dk/services/BepiPred-2.0/) to predict B-cell epitopes based on amino acid properties and immunological data (Peters, Nielsen, & Sette, 2020; Vita et al., 2015).

For inclusion, epitopes were selected based on several immunological and practical criteria: (i) high prediction scores above the recommended threshold, (ii) surface accessibility, (iii) appropriate length (typically 8–20 amino acids), and (iv) location in functionally relevant or conserved regions of the envelope protein. We also prioritized epitopes with predicted antigenicity, non-allergenicity, and non-toxicity to ensure safety and effectiveness.

Exclusion criteria included (i) epitopes located in highly variable or non-surface regions, (ii) low-scoring predictions, (iii) sequences overlapping with known host proteins to reduce the risk of autoimmunity, and (iv) those with potential allergenic or toxic properties based on downstream immunoinformatics screening.

This strategic selection ensured that only the most promising and biologically relevant epitopes were shortlisted for further processing in the multi-epitope vaccine design.

### Screening for Antigenic Potential, Allergenicity, and Toxicity

To ensure the immunological suitability and safety of the selected HIV-2 env protein epitopes, a thorough screening process was conducted focusing on their antigenic potential, allergenic profile, and toxicity. Antigenicity prediction was performed using the VaxiJen v2.0 (available at: https[:]//www[.]ddg-pharmfac[.]net/vaxijen/VaxiJen/VaxiJen.html) platform, a well-established server that relies on auto-cross covariance (ACC) transformation of protein sequences. A threshold of 0.45 was applied to discriminate likely antigens from non-antigenic peptides, prioritizing candidates with a higher potential to elicit immune activation.

To mitigate the risk of eliciting allergic reactions, each epitope was evaluated using AllerTOP v2.1 (available at: https[:]//www[.]ddg-pharmfac[.]net/allertop_test/), which classifies proteins based on physicochemical properties and alignment-free algorithms. Epitopes predicted as non-allergenic were favored to enhance the tolerability of the vaccine construct. Further, toxicity screening was carried out using ToxinPred (available at: http[:]//crdd[.]osdd[.]net/raghava/toxinpred/), a tool designed to identify toxic motifs in peptide sequences through a combination of SVM-based models and motif search.

Only those epitopes that fulfilled all three criteria—demonstrating antigenicity, non-allergenicity, and non-toxicity—were retained for the design of the multi-epitope vaccine construct. This multi-parameter screening strategy ensured the selection of immune-responsive yet biologically safe candidates for downstream vaccine development.

### Immuno-activation Potential of Epitopes for Cytokine Induction

To evaluate the immune-modulatory potential of the selected Major Histocompatibility Complex (MHC)-II epitopes, their ability to induce cytokine production was assessed using specialized prediction tools. The capacity of epitopes to stimulate interleukin-10 (IL-10) secretion was evaluated using the IL10Pred server (accessible at: https[:]//webs[.]iiitd[.]edu[.]in/raghava/il10pred/), while IL-4 induction potential was predicted with the IL4Pred tool (accessible at: https[:]//webs[.]iiitd[.]edu[.]in/raghava/il4pred/design.php). In addition, both MHC-I and MHC-II epitopes were under the observation to check the compatibility to induce interferon-gamma (IFN-γ), a very critical cytokine in antiviral responses, using the IFNepitope prediction server (accessible at: https[:]//webs[.]iiitd[.]edu[.]in/raghava/ifnepitope/predict.php). These predictions aids and ensure the selected epitopes must possess immunostimulatory properties needed for effective vaccine design.

### B-cell Epitope Evaluation and Antibody Induction Potential

The selected B-cell epitopes can stimulate an antibody-mediated immune response, we utilized the IgPred server (available at: https[:]//webs[.]iiitd[.]edu[.]in/raghava/igpred/). This tool allowed us to check the likelihood that each epitope could trigger immunoglobulin production, thereby confirming their role in activating the humoral immune system.

### Conservation Analysis of Epitopes

we execute a conservation analysis using the IEDB Epitope Conservancy Tool (accessible at: https[:]//tools[.]iedb[.]org/conservancy/). This analysis helped us determine the degree of sequence similarity between the chosen epitopes and the full-length protein sequence, ensuring that the epitopes are well-conserved and less prone to mutation-driven immune evasion.

### Assessment of HLA Distribution Across Populations

Selected epitopes must elicit immune responses in genetically diverse populations in multi-epitope vaccines. This is important because the human leukocyte antigen (HLA) system, which presents epitopes to T cells, is highly polymorphic and allele distributions vary by ethnicity and region. A detailed IEDB Population Coverage Tool analysis (available at: https[:]//tools[.]iedb[.]org/population/). solved this problem and maximized vaccine applicability. Using region-specific HLA allele frequency data and predicted epitope binding affinities, this tool estimates the percentage of a population likely to respond to at least one epitope.

We examined West African, Pakistani, and global populations. West Africa was included due to its high burden of endemic infectious diseases and HLA allele genetic diversity; Pakistan was selected for regional relevance given our study’s geographical focus; and global coverage was assessed to assess vaccine universality. We used IEDB-recommended algorithms NetMHCpan and NetMHCIIpan to predict strong-binding MHC-I & II epitopes. The population coverage calculations only used epitopes with IC50s below 500 nM and high predicted binding affinity. Epitope peptides and HLA molecules must interact strongly for antigen presentation and T cell activation.

Key parameters were calculated by IEDB Population Coverage Tool. The HLA genotype-based epitope set response percentage of each region’s population was estimated. Population coverage (%) estimates vaccination response. To measure population immune response breadth, the tool calculated the average epitope-HLA combinations recognized per person. A higher average suggests a stronger, more complex immune response, improving vaccine efficacy even with poorly processed epitopes. The minimum epitope-HLA interactions needed to cover 90% of the population, PC90, assessed vaccine inclusivity and public health impact.

Also important in epitope selection was conservation.We tested each predicted epitope for sequence conservation across pathogen strains or isolates using IEDB’s epitope conservancy analysis tool. In circulating strains, high-conservancy epitopes are immunologically relevant due to their low mutation rate. Thus, immune escape is reduced and the vaccine candidate works.

We identified a subset of epitopes with high immunogenic potential and expected to elicit responses in a large proportion of the global population, including underrepresented West African and South Asian groups, using HLA allele frequency, epitope binding affinity, and sequence conservancy data. A comprehensive approach improves the vaccine’s translational relevance and clinical utility in various clinical settings.

### Peptide-Based Vaccine Construction Targeting Multiple Epitopes

For the assembly of the vaccine candidate, epitopes exhibiting strong binding affinity and immunogenic potential—spanning B-cell, MHC-I, & MHC-II categories—from the HIV-2 env. protein was carefully chosen. These selected epitopes were concatenated using the flexible CPGPG linker sequence, which facilitates proper folding and optimal exposure of individual epitopes to the immune system, as supported by previous studies (Dey et al., 2022).

To augment the overall immunogenic profile of the vaccine, the HP91 peptide adjuvant was incorporated at the N-terminal region through an EAAAK rigid linker. This design choice increase antigen presentation and promotes activation of innate immune receptors, particularly Toll-like receptors (TLRs), thereby potentiating adaptive immune responses (Pulendran, Arunachalam, & O’Hagan, 2021). The integration of linkers and adjuvants not only contributes to the structural stability of the multi-epitope construct but also maximizes its ability to stimulate targeted immune mechanisms.

### Evaluation of Physicochemical Properties of the Vaccine Construct

The ProtParam tool (available at: https[:]//web[.]expasy[.]org/protparam/) was used to evaluate many physicochemical parameters of the engineered vaccine construct. Parameters such as molecular weight, theoretical isoelectric point (pI), aliphatic index, instability index GRAVY score, and predicted half-life in both Escherichia coli and mammalian cell environments were examined to anticipate the vaccine’s biochemical behavior. The GRAVY score, giving the overall hydrophobic or hydrophilic nature of the protein, provides insight into its solubility characteristics, which impact the formulation strategies and feasibility (Gasteiger et al., 2005). Notably, the molecular weight of the designed multi-epitope construct fell within the accepted range for peptide-based vaccines, approximately 30 to 110 kilodaltons, suitability for effective production and delivery.

### Solubility Analysis

we performed computational solubility prediction using two important validated computational tools: Protein-Sol (available at: https[:]//protein-sol[.]manchester[.]ac[.]uk). & the SCRATCH Protein Projection tool (available at: https[:]//scratch[.]proteomics[.]ics[.]uci[.]edu) (Hebditch et al., 2017; Magnan et al., 2009). Commonly used in recombinant protein synthesis, these tools project the probability of the protein remaining soluble when overexpressed in a bacterial host such Escherichia coli. Vaccine design depends much on solubility since it directly affects the efficiency of protein expression, purification, formulation, and downstream handling. Additionally, these tools also provide half-life and stability predictions, offering insights into the protein’s behavior under physiological conditions. Constructs predicted to have high solubility and favorable stability profiles were considered more suitable for therapeutic development and large-scale manufacturing.

### Secondary Structure Analysis

To assess the secondary structural composition of the vaccine construct, computational predictions were performed using two widely validated bioinformatics tools: SOPMA (Self-Optimized Prediction Method with Alignment) and PSIPRED. SOPMA (accessible at: https[:]//npsa-prabi[.]ibcp[.]fr/cgi-bin/npsa_automat.pl?page=/NPSA/npsa_sopma.htm) predicts structural motifs by aligning query sequences with known protein structures, while PSIPRED (accessible at: https[:]//bioinf[.]cs[.]ucl[.]ac[.]uk/psipred/) applies neural network-based learning to forecast folding patterns from primary amino acid sequences (Geourjon & Deleage, 1995; McGuffin, Bryson, & Jones, 2000).

Both tools were used to identify the distribution of alpha-helices, beta-turns, extended strands and random coils which are key indicators of the protein’s folding and functional potential. SOPMA and PSIPRED have demonstrated predictive accuracies of approximately 80% and 78.1%, respectively, making them reliable for early-stage structural modeling. The results provided foundational insights into the vaccine’s conformational characteristics, essential for downstream modeling and functional analysis.

### Computational Tertiary Structure Modeling

The three-dimensional (tertiary) structure of the designed vaccine construct was modeled using the 3DPRO module, part of the SCRATCH Protein Predictor suite (available at: https[:]//scratch[.]proteomics[.]ics[.]uci[.]edu). This tool utilizes machine learning-based algorithms that integrate sequence-derived features and generate a predicted 3D conformation of the input protein sequence (Cheng, Randall, Sweredoski, & Baldi, 2005). By translating primary sequence data into spatial structural information, 3DPRO predicts protein’s potential folding patterns, surface exposure, and interaction capabilities, which are critical for evaluating antigen presentation, stability & receptor binding.

### Protein Simulation and Structure Validation

To confirm the structural integrity and suitability of the recombinant protein as a vaccine candidate. The predicted tertiary structure was visualized and saved in PDB format for detailed analysis. We used PyMOL 3.1 (available at: https[:]//www[.]pymol[.]org) to examine the 3D conformation, allowing us to assess critical structural features such as folding, surface accessibility, and potential epitope exposure. This step ensured that the protein model retained a stable and biologically relevant structure, which is essential for effective immune recognition and vaccine efficacy.

### Validation of the Predicted 3D Structure

The predicted 3D structure of the multi-epitope protein was validated using multiple tools to assess its accuracy and structural quality. ProSA (!!https://prosa.services.came.sbg.ac.at/prosa.php) was used to evaluate overall model quality based on atomic interactions, while PROCHECK (available at: https[:]//www[.]ebi[.]ac[.]uk/thornton-srv/software/PROCHECK/) generated Ramachandran plots to analyze stereochemical properties. Additional phi (ϕ) and psi (ψ) angle assessments were performed using the RamachandranPlot Server (accessible at: https[:]//zlab[.]umassmed[.]edu/bu/rama/). The Verify3D module of SAVES v6.0 (available at: https[:]//servicesn[.]mbi[.]ucla[.]edu/SAVES/) was employed to assess the compatibility between the 3D model and its amino acid sequence. Finally, GalaxyRefine (available at: https[:]//galaxy[.]seoklab[.]org/cgi-bin/submit.cgi?type=REFINE) was used for structure refinement and quality improvement based on multiple structural parameters.

### Prediction of Conformational B Cell Epitopes

Usually in PDB form, conformational B-cell epitopes are identified by means of three-dimensional protein structure analysis, so enabling surface-exposed areas most likely to be recognized by antibodies. To forecast these epitopes, we used the ElliPro tool housed on the IEDB web server (accessible at: https[:]//tools[.]iedb[.]org/ellipro/). Using the Protrusion Index (PI), which measures the solvent accessibility of residues, ElliPro assesses the structure of the protein highlighting those most likely to interact with B-cell receptors. This method allows correct mapping of discontinuous epitopes vital for humoral immunity (Ponomarenko et al., 2008).

### Molecular Docking of Vaccine Construct with Toll-Like Receptors

For the evaluation of potential immunostimulatory interactions between designed vaccine construct and components of the innate immune system, molecular docking analyses were performed with selected Toll-Like Receptors (TLRs). The ClusPro 2.0 web server (accessible at: https[:]//cluspro[.]org) was used to perform rigid-body protein–protein docking. This platform utilizes the PIPER algorithm, which applies a Fast Fourier Transform (FFT)-based correlation approach to predict optimal binding conformations.

Structural data for TLR targets were obtained from the Protein Data Bank, using the following PDB IDs: 2AOZ, 3FXL, 3WPB, and 1Z1W. The resulting vaccine and TLR complexes were analyzed with PDBsum tool (available at: https[:]//www[.]ebi[.]ac[.]uk/thornton-srv/databases/pdbsum/) which provides hydrogen bonds, salt bridges, and hydrophobic interactions. This docking study aimed to see the binding affinity and interaction patterns between the vaccine construct and innate immune receptors, supporting its potential immunogenic performance in vivo.

### Evaluating Structural Flexibility through Normal-Mode Analysis

The normal mode analysis (NMA) was carried out by using the iMODS web server (accessible at: http[:]//imods[.]chaconlab[.]org) (Lopéz-Blanco, Garzón, & Chacón, 2011). The NMA give us the intrinsic flexibility of macromolecular structures by simulating their collective motions based on internal coordinates.

iMODS help characterize how a protein complex behaves under physiological conditions. These include deformability plots, B-factor estimations (which mirror atomic fluctuations), eigenvalues (reflecting overall motion stiffness), and covariance matrices that capture both coordinated and opposing movements between residues.

### Molecular Dynamics Simulation for Assessing Vaccine–TLR2 Complex Stability

Molecular Dynamics (MD) simulations with GROMACS version 2021.5 (available at: https[:]//manual[.]gromacs[.]org/2021.5/download.html) give us the structural integrity and dynamic behavior of the best vaccination TLR2 complex. To avoid edge effects, the protein– protein complex was solvated in a TIP3P water model in a cubic simulation box with a 1 nm buffer zone. The CHARMM36 force field (Vanommeslaeghe et al., 2010) generated system topologies. Counter-ions (Na⁺ and Cl⁻) were added to achieve system neutrality.

Two phases of equilibration followed steepest descent energy minimization. Starting with a modified Berendsen thermostat, an NVT ensemble was run for 1 ns at 300 K. A Berendsen barostat-controlled 1 ns NPT ensemble at 1 bar (constant number of particles, pressure, and temperature) followed (Berendsen et al., 1984). A 100-ns production run under periodic boundary conditions replicated the complex’s behavior in a biologically plausible environment.

With a 1.0 nm cutoff (Petersen, 1995), the Particle Mesh Ewald (PME) approach handled long-range electrostatic interactions to ensure correct charge distribution in the simulation. Post-simulation analysis revealed the vaccine–receptor complex’s stability, flexibility, and compactness over time by focusing on RMSD, RMSF, and Rg.

### Disulfide Bond Prediction

For enhancing stability of the vaccine potential disulfide-bonds were projected using Disulfide by using a tool named as Design 2 (DbD2) (available at: http[:]//cptweb[dot]cpt[dot]wayne[dot]edu/DbD2/) This tool identifies suitable cysteine residue pairs for mutation by analyzing their spatial proximity and geometric orientation. Candidate mutations with favorable χ3 dihedral angles between −87° and +97° and energy scores below 2.2 kcal/mol were considered optimal for forming stable disulfide bridges, which can improve protein folding and structural rigidity (Craig & Dombkowski, 2013).

### Simulation of Host Immunogenicity Against the Vaccine Construct

To gain a predictive understanding of how the vaccine construct might interact with the human immune system, we used C-ImmSim (available at: !!https[:]//kraken.iac.rm.cnr.it/C-IMMSIM), a powerful agent-based simulation platform that mimics complex immunological processes in silico. This tool uses position-specific scoring matrices and immune network theory to model immune cell activation, expansion, and regulation. It includes B lymphocytes, CD4⁺ helper T cells, CD8⁺ T lymphocytes (CTL), and antibody-secreting plasma cells.

These parameters aid in predicting the potential strength, duration, and specificity of the immune reaction show by the candidate vaccine we look in the expected real-world performance (Rapin et al., 2010; Ragone et al., 2021).

### Optimizing Codon Usage for Enhanced Protein Expression

The efficient production of the vaccine in choose host system, the DNA sequence of the vaccine construct was optimized with using Java Codon Adaptation Tool (JCat) (available at: https[:]//www[.]jcat[.]de).

This optimization describes the original amino acid sequence while improving the chances of successful recombinant expression that is an essential step for reliable and vaccine manufacturing (Grote et al., 2005).

### Expression Vector Selection

A desired bacterial expression vector was selected from the Helmholtz Munich Vector Database (available at: https[:]//www[.]helmholtz-munich[.]de/pepf/materials/vector-database/bacterialexpression-vectors/) based on its compatibility with downstream experimental requirements as well as protein expression parameters (Crass et al., 2004).

### In Silico Cloning and Vector Validation

Serial Cloner v2.1 (available at: https[:]//serial-cloner[.]en[.]softonic[.]com) was used to simulate the cloning process and made sure that the vaccine construct was properly included. This tool made it easier to identify appropriate enzyme restriction sites and alss ensures insertion of codon-optimized vaccine sequence.

The optimized gene was cloned in silico into the bacterial expression vector pET28a(+), allowing us to confirm the proper orientation and compatibility for efficient protein expression in a prokaryotic host system (Humayun et al., 2022). This step serves as a crucial checkpoint before proceeding to laboratory-based cloning and expression.

## RESULTS

### HIV-2 env Protein Sequences

The reference amino acid sequence of the HIV-2 *env* polyprotein was retrieved from the NCBI database for downstream epitope mapping and vaccine design. This sequence served as the primary input for B-cell and T-cell epitope prediction, antigenicity profiling, and structural modeling in subsequent *in silico* analyses.

MMNQLLIAILLASACLVYCTQYVTVFYGVPTWKNATIPLFCATRNRDTWGTIQCLPDNDDYQEITLNVTEAFDAWNNTVTEQAIEDVWHLFETSIKPCVKLTPLCVAMKCSSTESSTGNNTTSKSTSTTTTTPTDQEQEISEDTPCARADNCSGLGEEETINCQFNMTGLERDKKKQYNETWYSKDVVCETNNSTNQTQCYMNHCNTSVITESCDKHYWDAIRFRYCAPPGYALLRCNDTNYSGFAPNCSKVVASTCTRMMETQTSTWFGFNGTRAENRTYIYWHGRDNRTIISLNKYYNLSLHCKRPGNKTVKQIMLMSGHVFHSHYQPINKRPRQAWCWFKGKWKDAMQEVKETLAKHPRYRGTNDTRNISFAAPGKGSDPEVAYMWTNCRGEFLYCNMTWFLNWIENKTHRNYAPCHIKQIINTWHKVGRNVYLPPREGELSCNSTVTSIIANIDWQNNNQTNITFSAEVAELYRLELGDYKLVEITPIGFAPTKEKRYSSAHGRHTRGVFVLGFLGFLATAGSAMGAASLTVSAQSRTLLAGIVQQQQQLLDVVKRQQELLRLTVWGTKNLQARVTAIEKYLQDQARLNSWGCAFRQVCHTTVPWVNDSLAPDWDNMTWQEWEKQVRYLEANISKSLEQAQIQQEKNMYELQKLNSWDIFGNWFDLTSWVKYIQYGVLIIVAVIALRIVIYVVQMLSRLRKGYRPVFSSPPGYIQQIHIHKDRGQPANEETEEDGGSNGGDRYWPWPIAYIHFLIRQLIRLLTRLYSICRDLLSRSFLTLQLIYQNLRDWLRLRTAFLQYGCEWIQEAFQAAARATRETLAGACRGLWRVLERIGRGILAVPRRIRQGAEIALL.

### Features of the Selected env Protein

The sequence was made up of 858 amino acids, and its molecular weight was determined to be 98,824.48 Da using calculations. It has been determined that the theoretical isoelectric point, also known as pI, is 8.73. On the other hand, the total number of positively charged residues (Arg + Lys) was 94, while the number of negatively charged residues (Asp + Glu) was 79. When these characteristics were taken into consideration, the protein was determined to be stable.

### Prediction of B and T Cell Epitopes

Linear B-cell epitopes predicted by BepiPred tool epitopes showed scores ranging from -0.006 to 2.441, with an average of -0.065, identifying regions likely to be recognized by antibodies.

T-cell epitope prediction was performed using NN-align 2.3 for MHC-II and NetMHCpan 4.1 EL for MHC-I alleles. High-affinity binders were selected based on percentile ranks and binding scores, and the most promising epitopes are listed in Tables 1, 2, and 3.

**Table 1.**
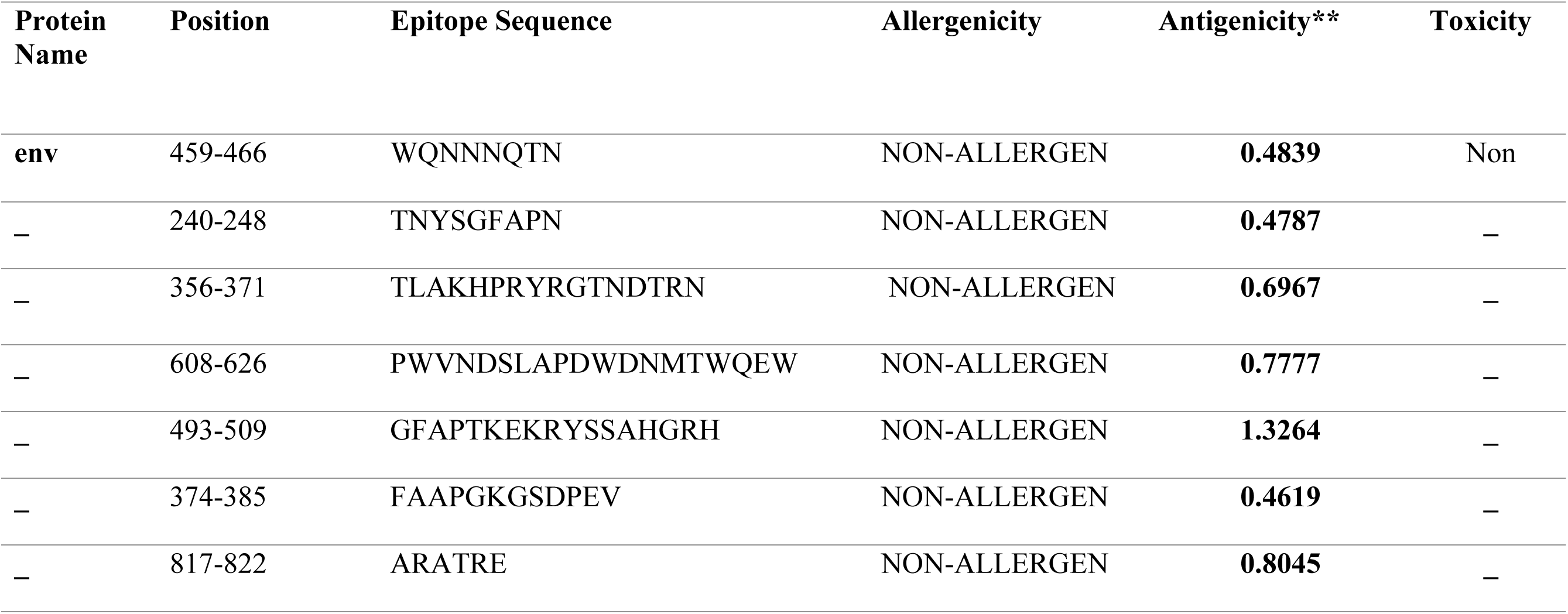
Bepipred Linear Epitope Prediction Results.

**Table 2.**
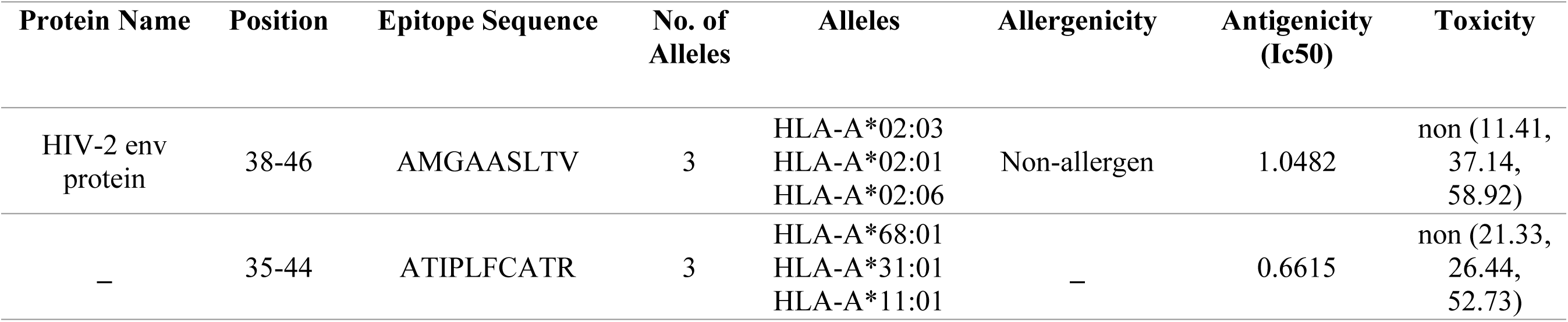
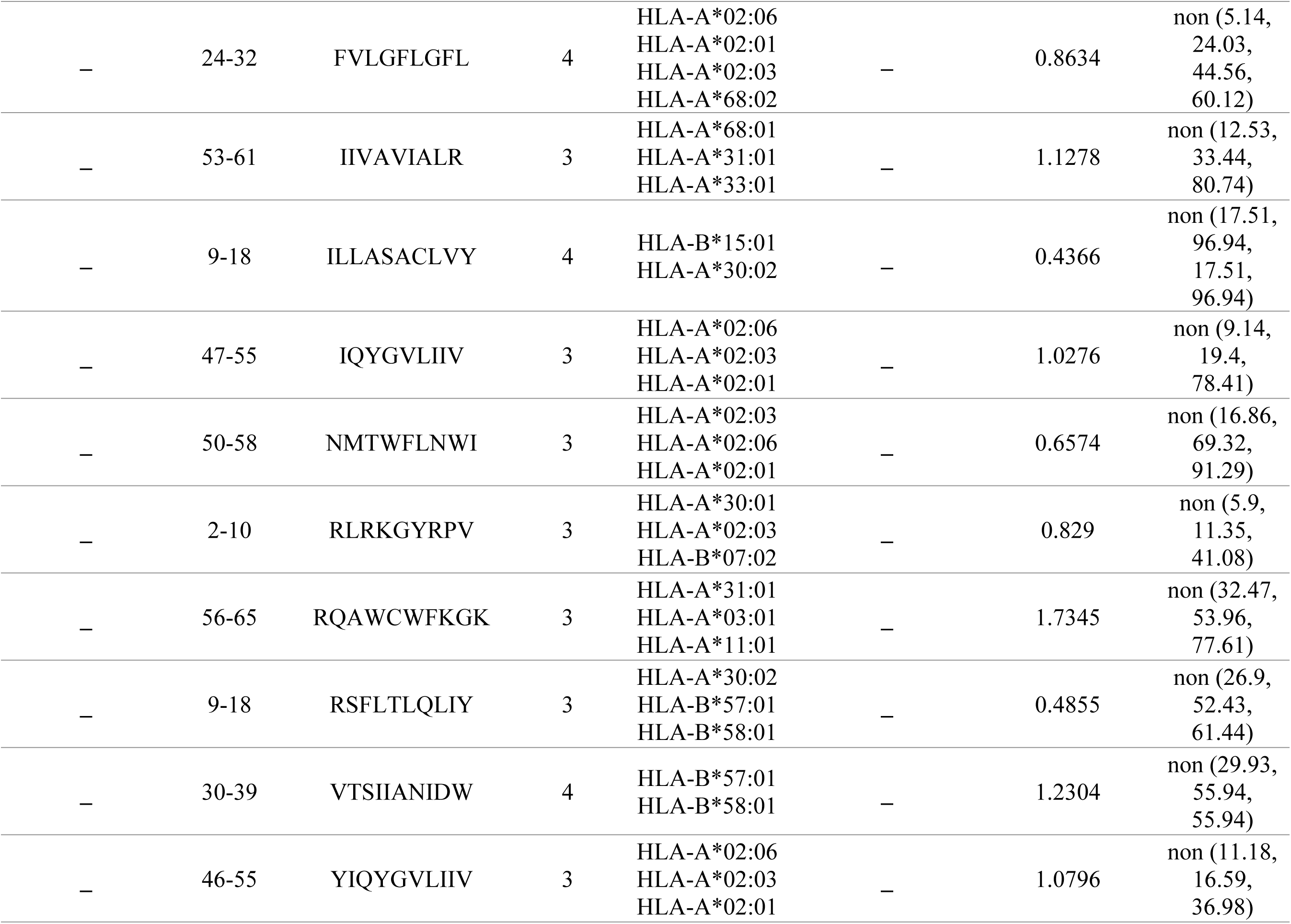
The selection of MCH 1 epitopes of HIV-2 env protein based on binding affinity.

### Cytokine Induction Potential of Epitopes

We evaluated selected MHC-II epitopes for their ability to induce IL-10 and IL-4 using predictive tools. Peptides such as LLSRSFLTLQLIYQN, SRSFLTLQLIYQNLR, RSFLTLQLIYQNLRD, PIAYIHFLIRQLIRL, and WPWPIAYIHFLIRQL showed high IL-10 induction scores. Similarly, PIAYIHFLIRQLIRL, LTSWVKYIQYGVLII, and their adjacent variants were strong IL-4 inducers. IFN-γ analysis confirmed that all MHC-I & MHC-II epitopes had positive predictions, with RSFLTLQLIYQNLRD, ILLASACLVY, and AMGAASLTV among the top candidates (Table 5, 6,10 and 11).

**Table 3.**
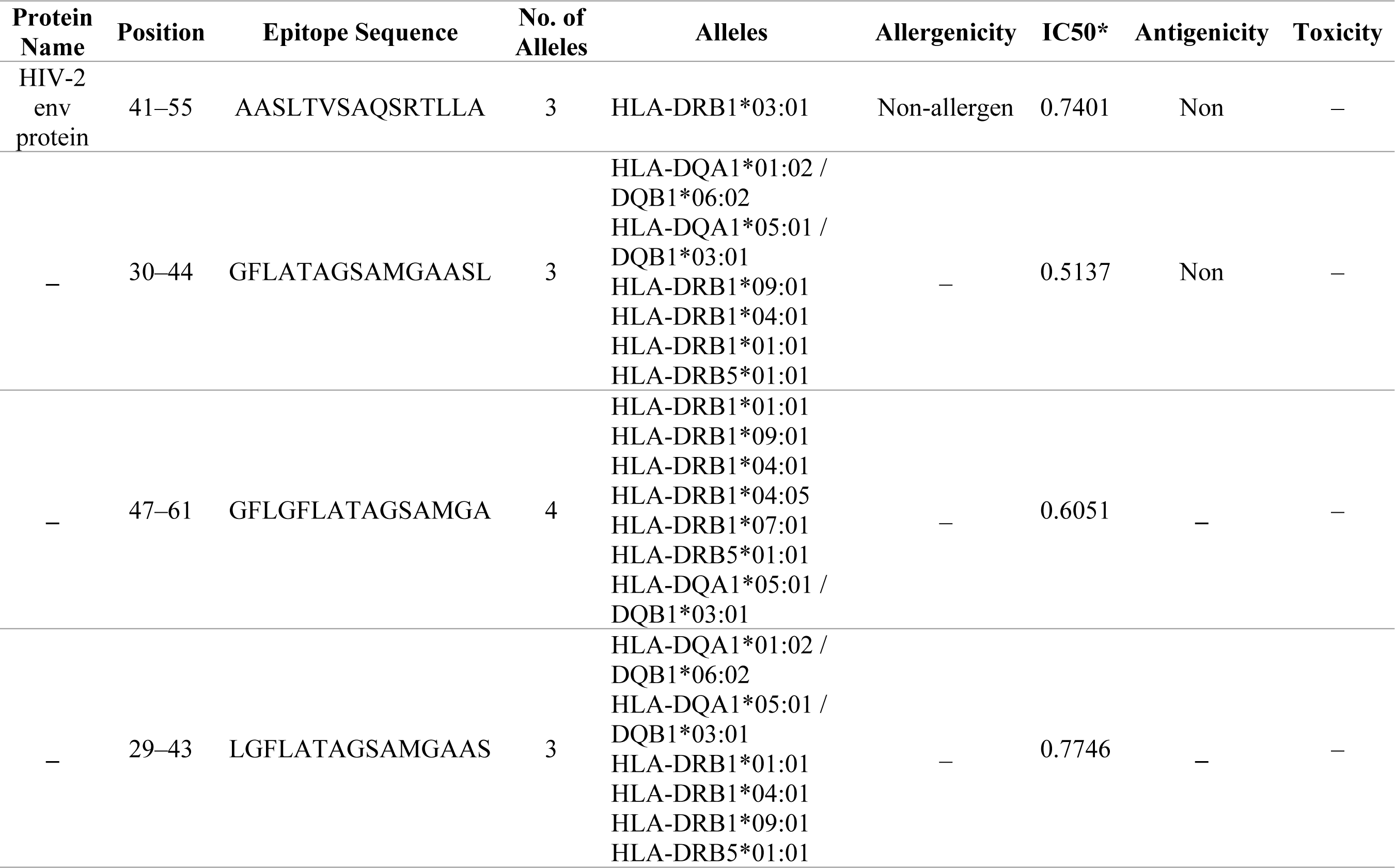
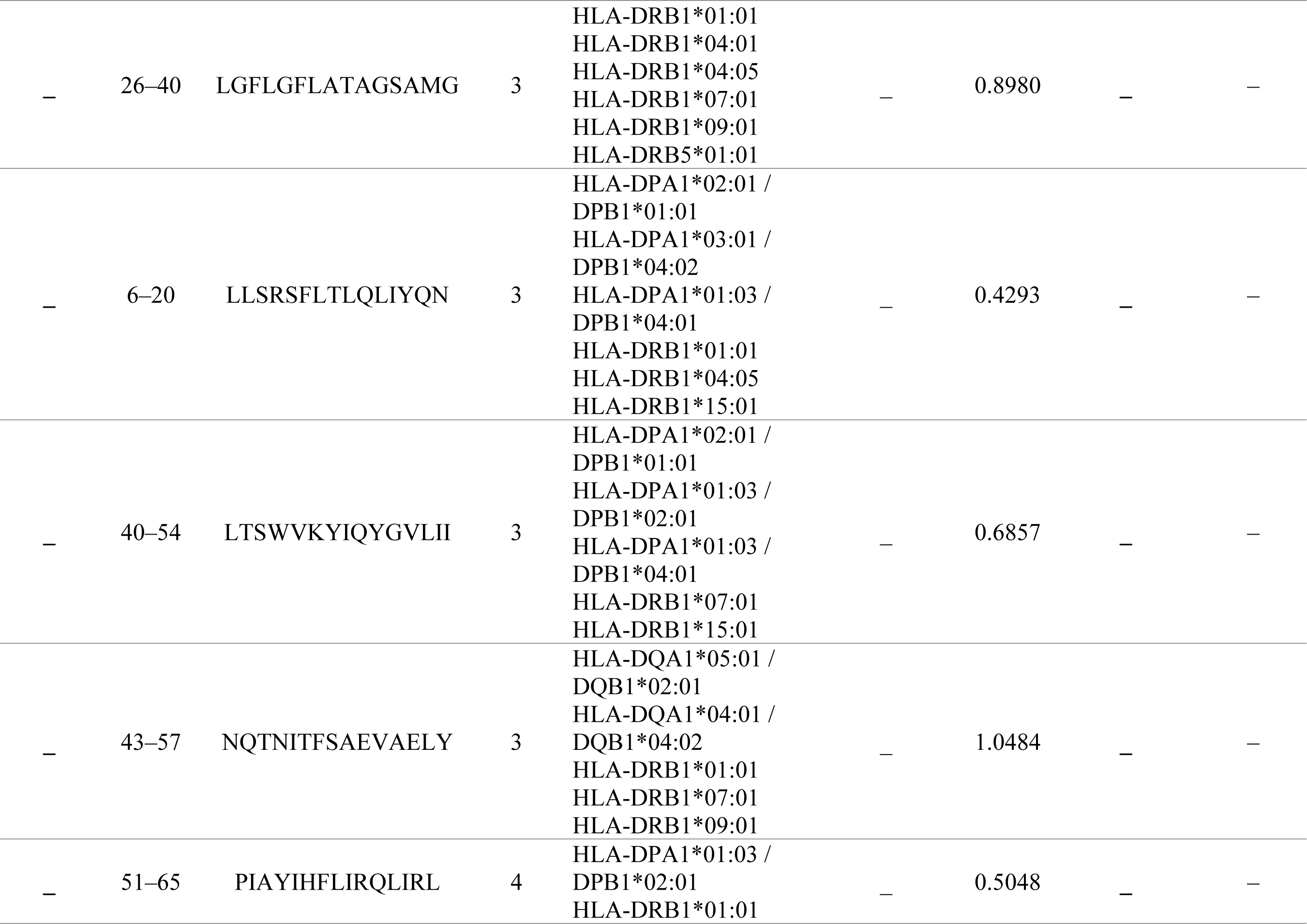
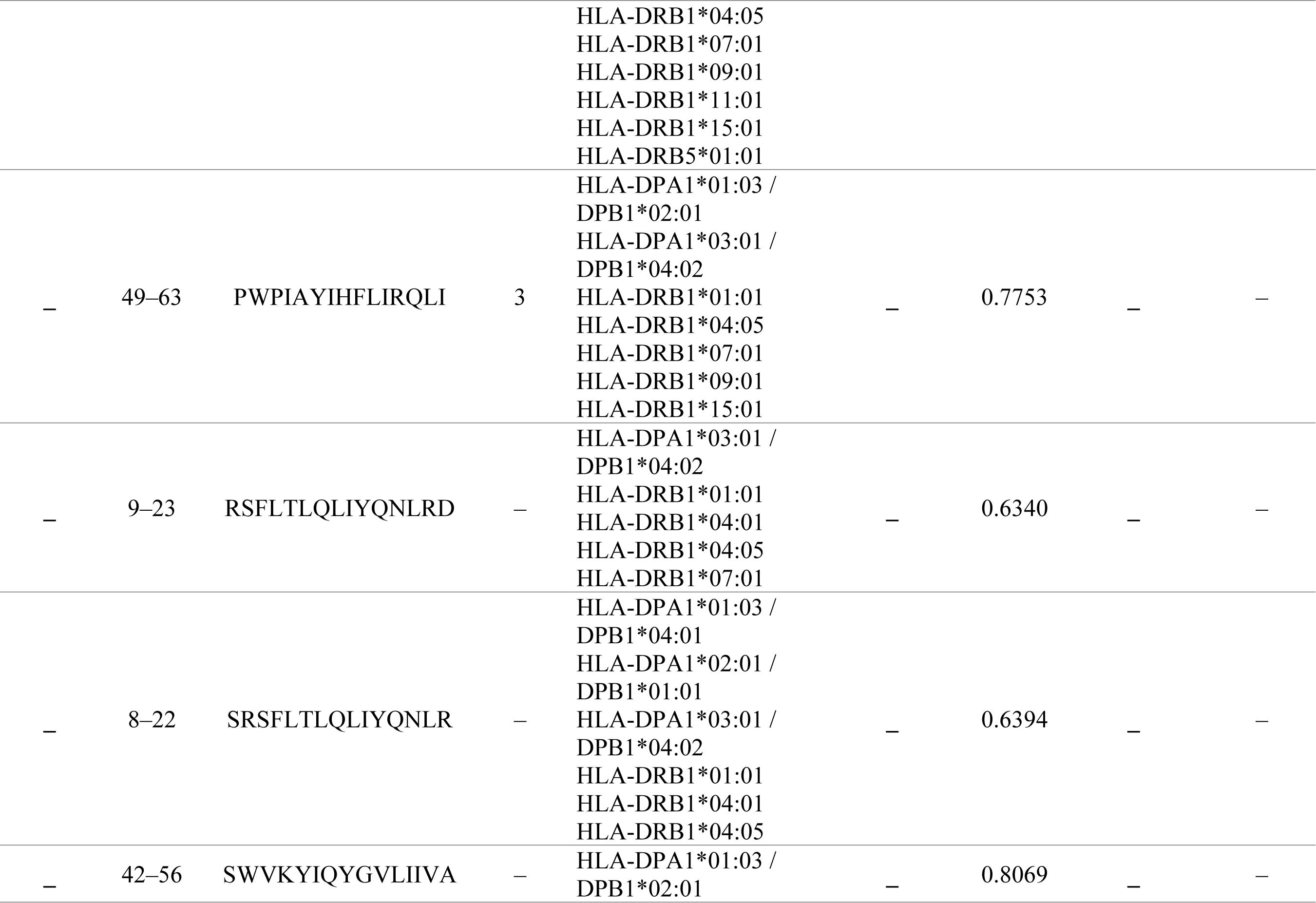
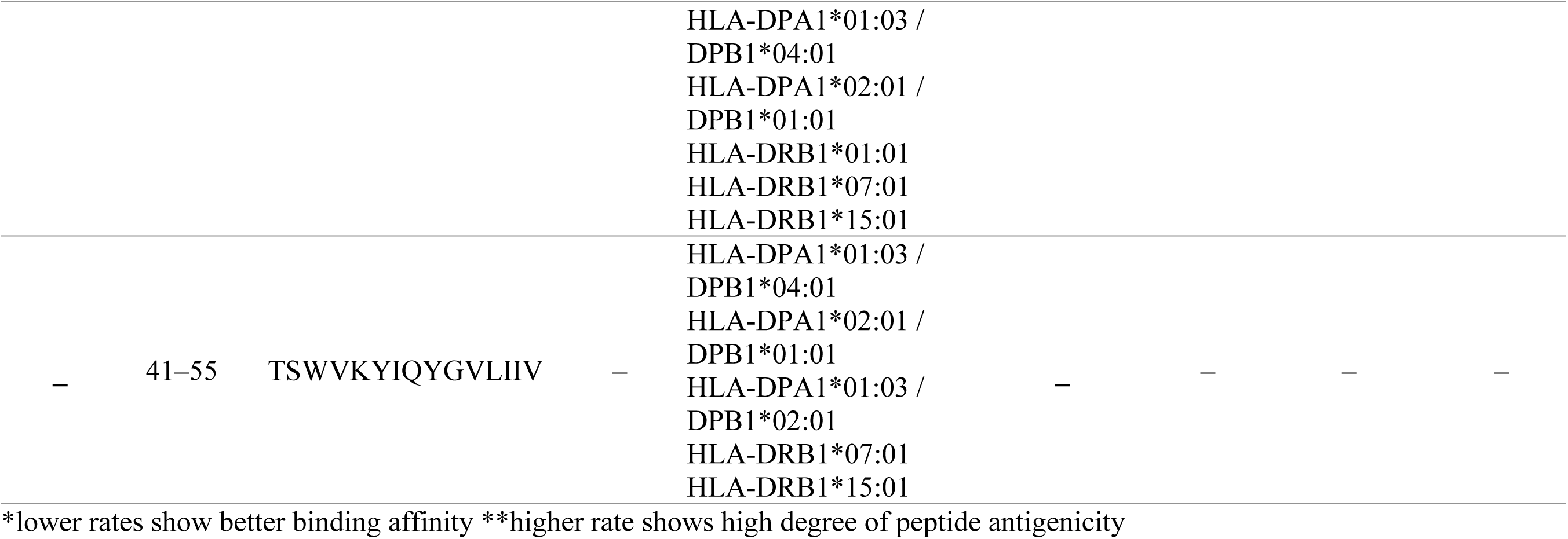
The selection of MCH 2 epitopes of HIV-2 env protein based on binding affinity.

**Table 4.**
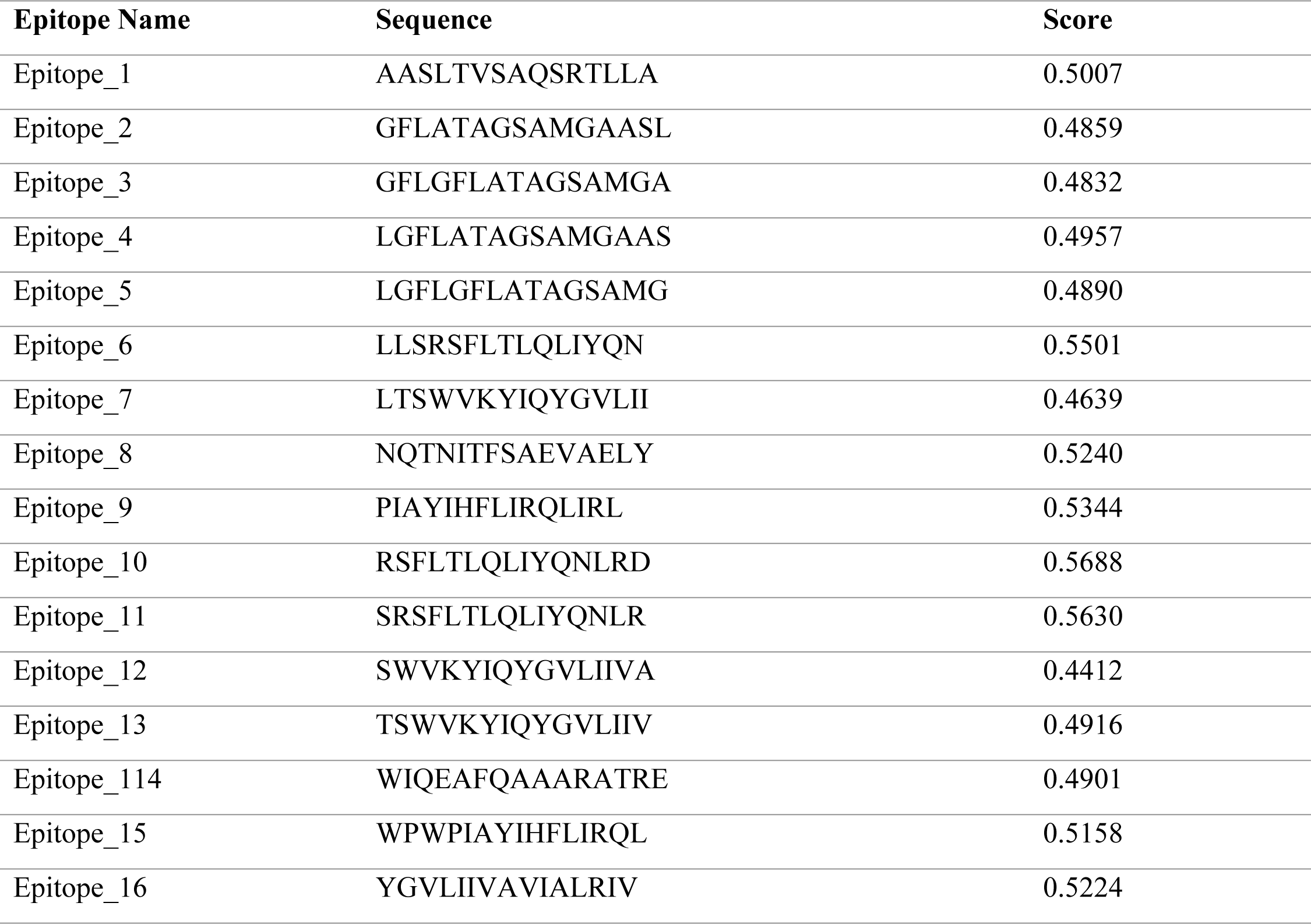
Positive SVM based Predicted IFN-γ Epitope for MHC-II (HTL) Candidates.

**Table 5.**
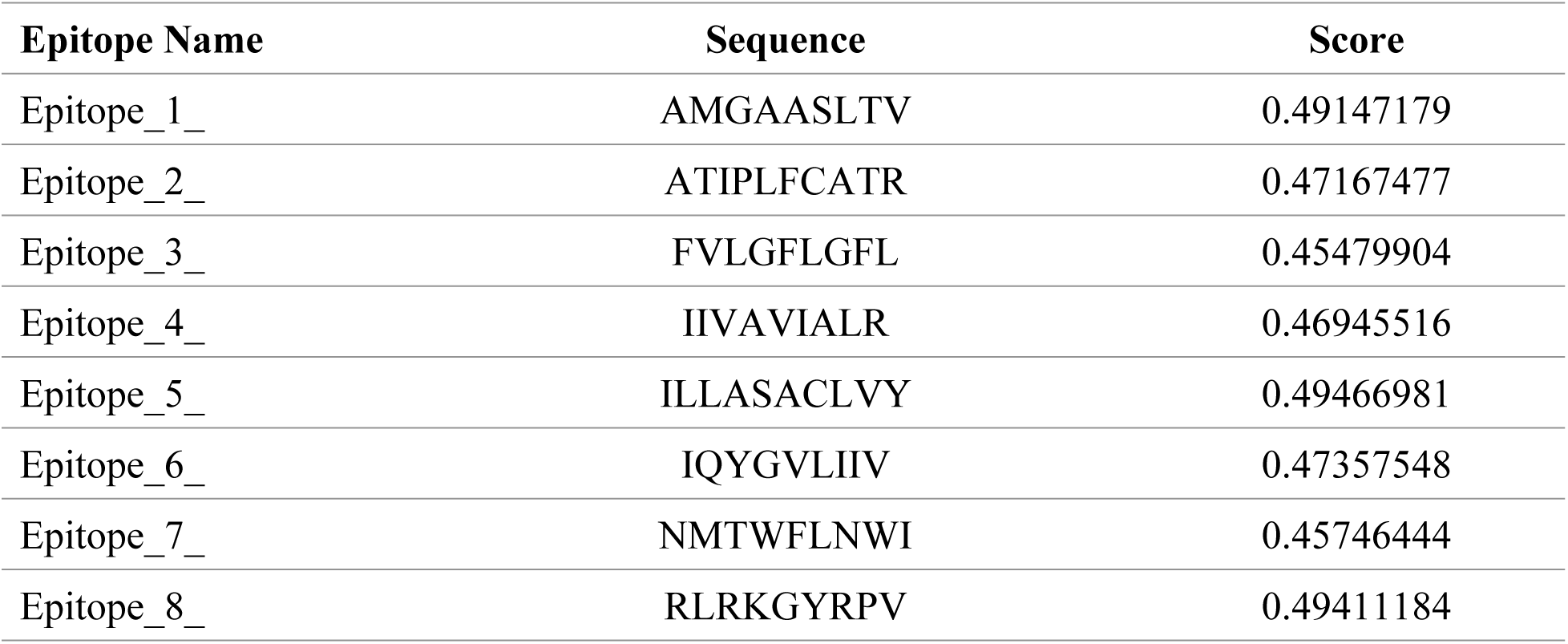
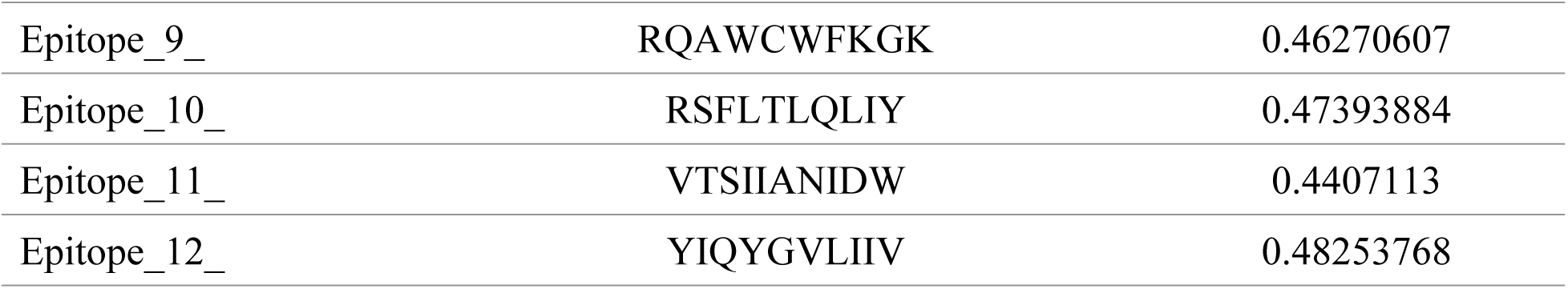
Positive SVM based Predicted IFN-γ Epitope for MHC-I (CTL) Candidates.

### Predicted B-cell Epitope and Antibody Response

Using IgPred, we assessed seven B-cell epitopes for their ability to elicit antibody responses. Only Epitope 5 (GFAPTKEKRYSSAHGRH) was strongly predicted to induce IgG (score: 0.925), suggesting potential for antibody-mediated immunity. The rest, including Epitope 3 (TLAKHPRYRGTNDTRN), fell below the positivity threshold and were considered non-epitopic (Table 9).

**Table 6.**
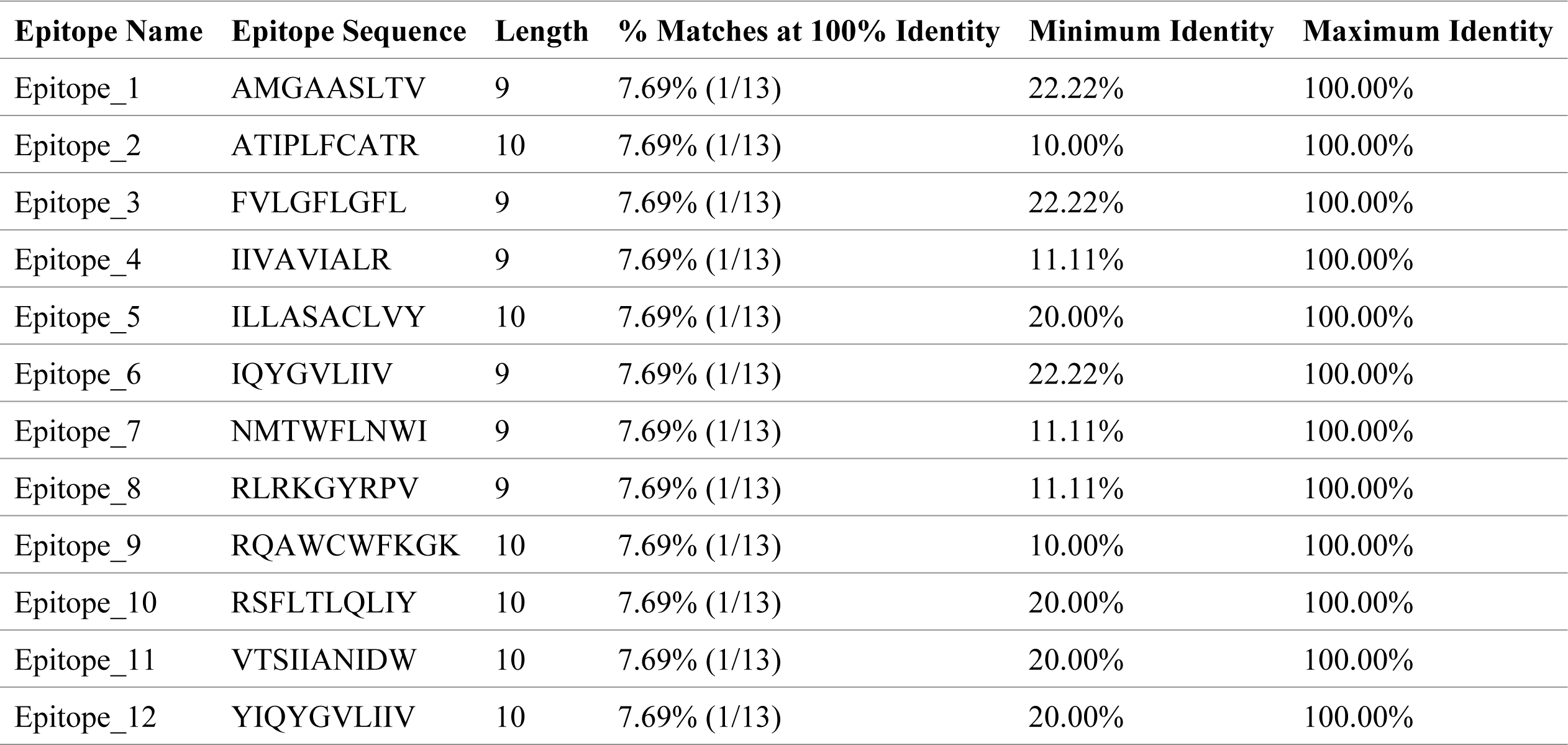
Conservancy analysis of predicted MHC-I epitopes.

**Table 7.**
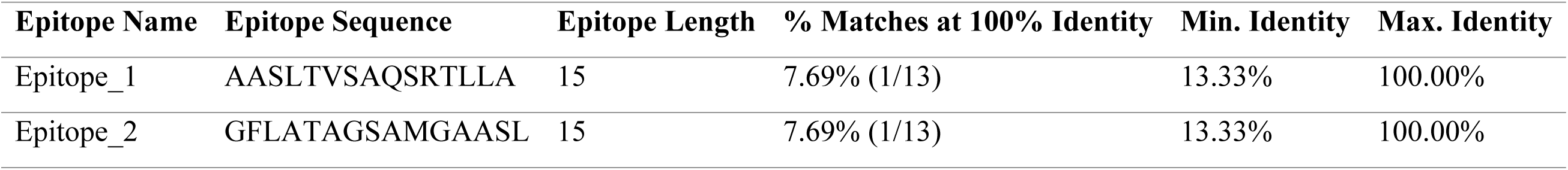
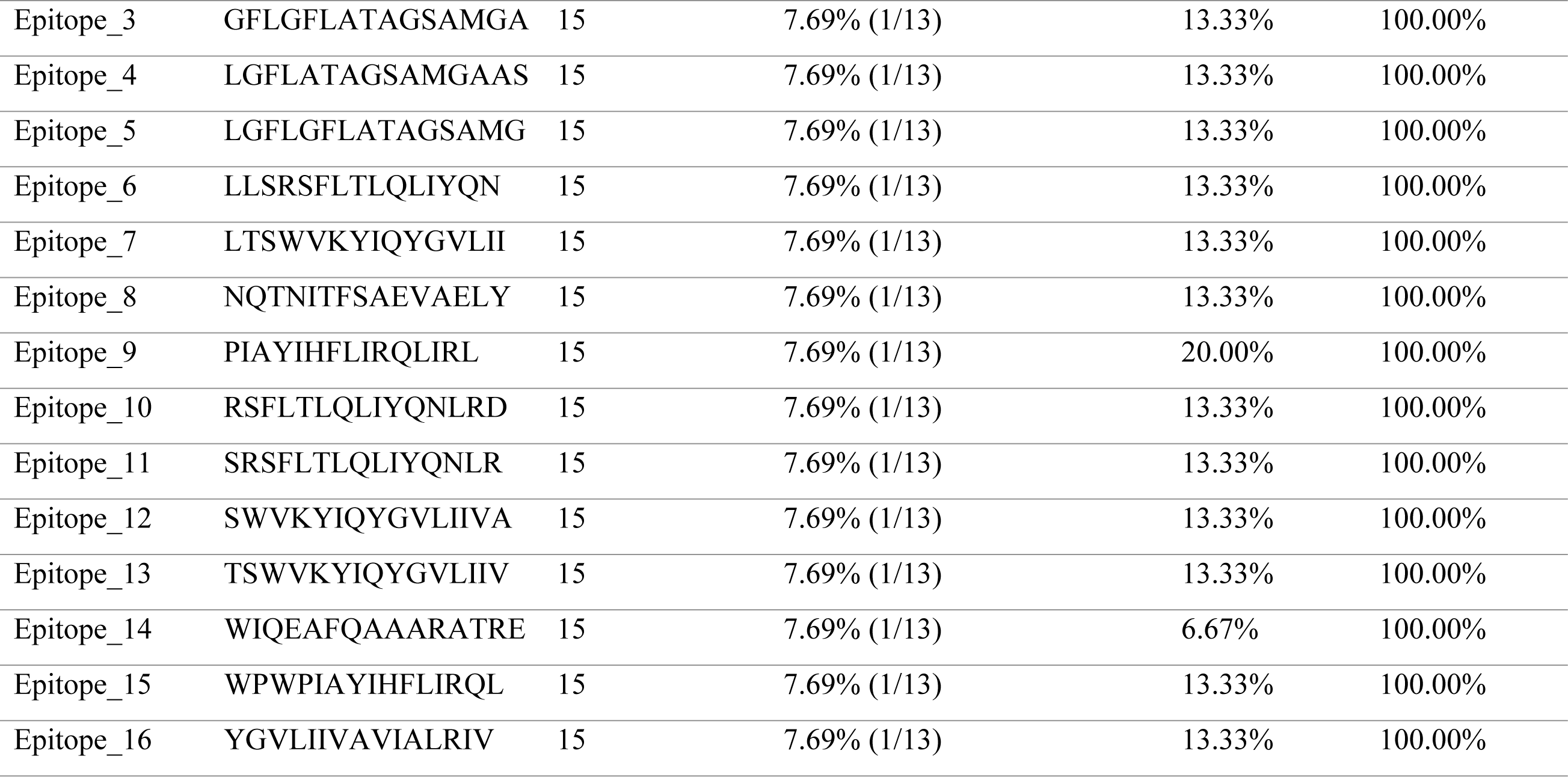
Epitope Conservancy Analysis for MHC-II.

**Table 8.**
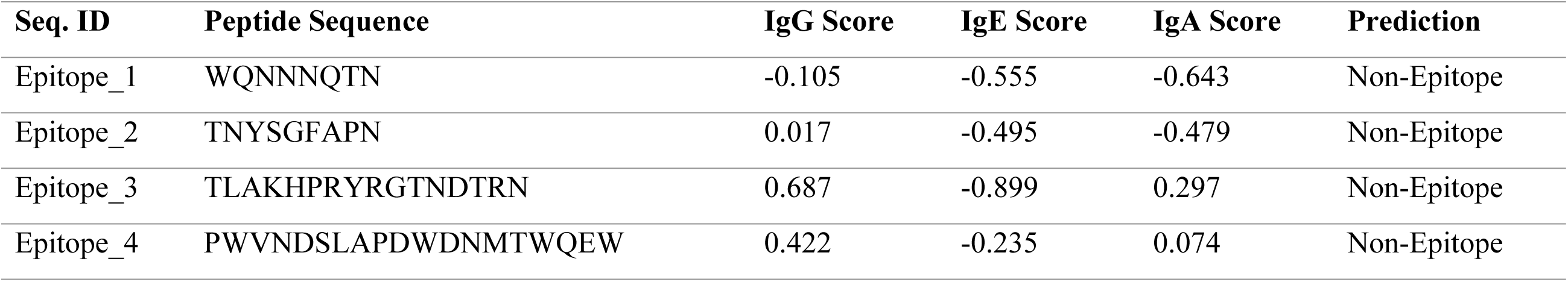
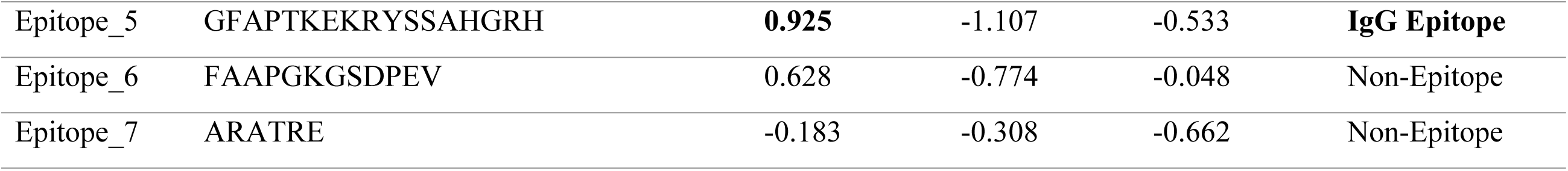
Antibody-specific B-cell epitope screening results.

**Table 9.**
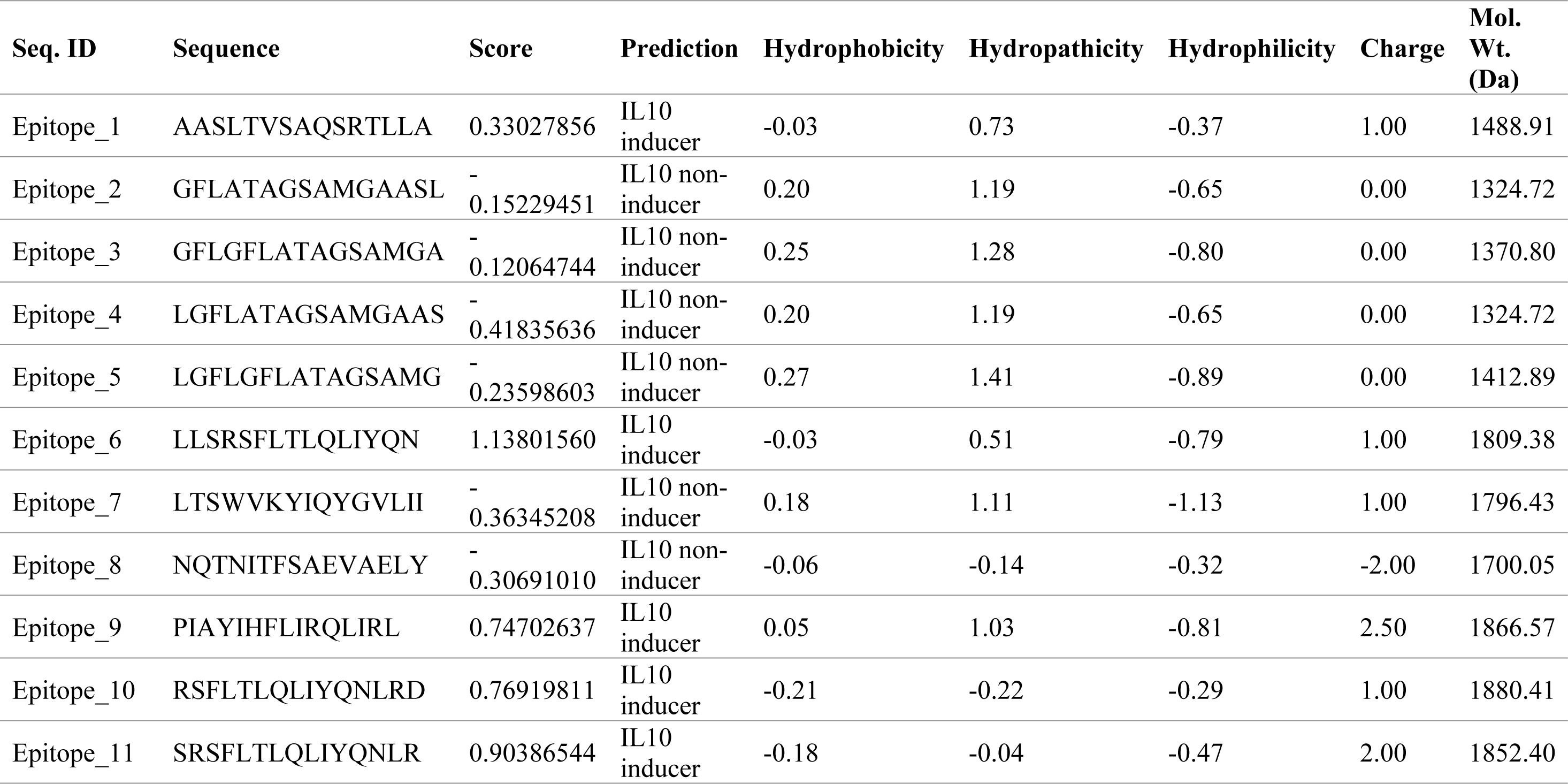
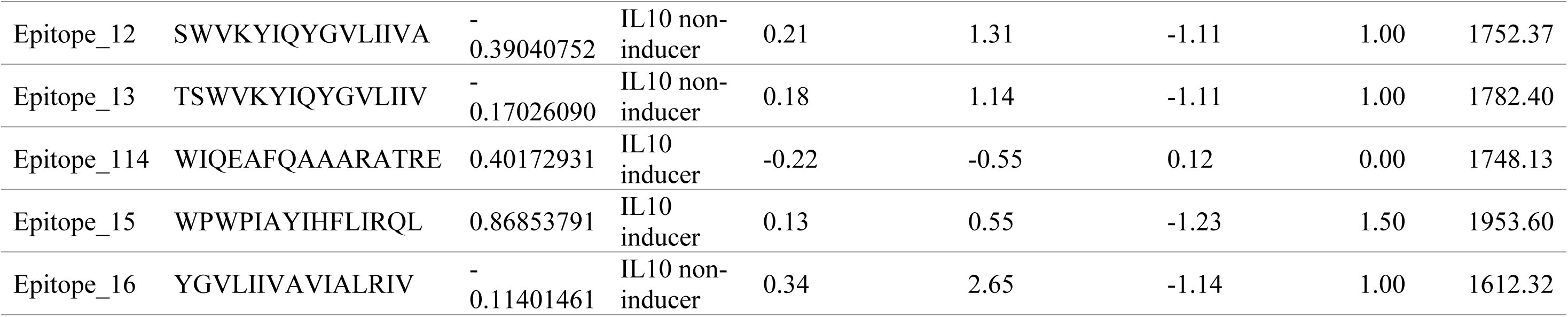
Prediction of IL-10 inducing epitopes from MHC-II candidates.

**Table 10.**
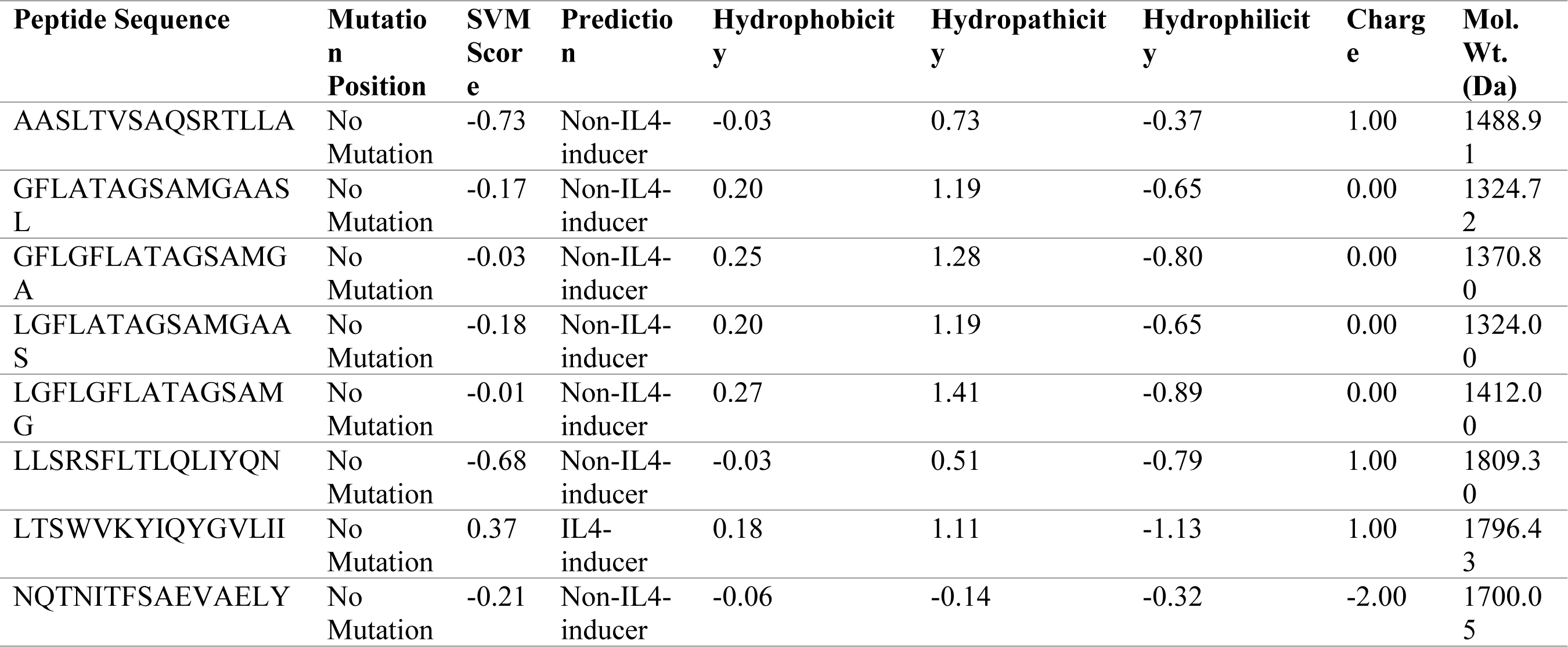
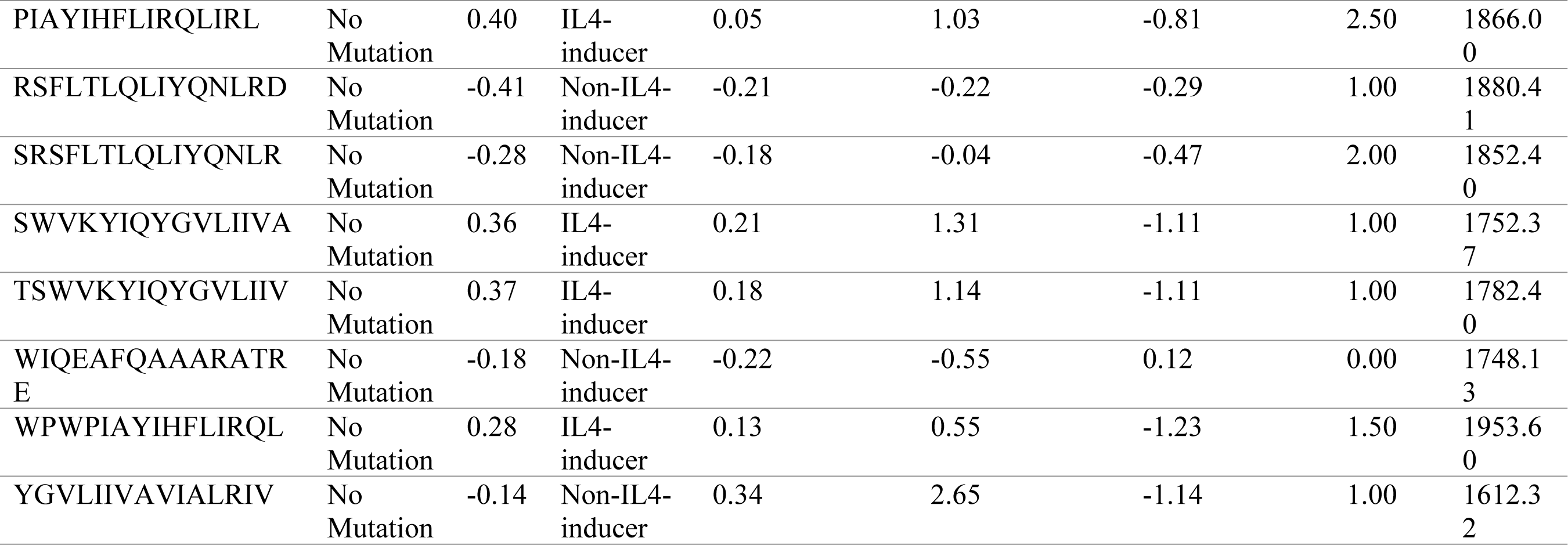
Prediction of IL-4 inducing epitopes among MHC-II candidates.

**Table 11.** Physicochemical Properties of Vaccine Construct.

| Characteristic | Details |
| --- | --- |
| Amino Acid Count | 477 residues |
| Calculated Molecular Mass | 53,279.18 Daltons |
| Predicted Isoelectric Point (pI) | 9.89 |
| Acidic Residue Content (Asp + Glu) | 18 residues |
| Basic Residue Content (Arg + Lys) | 39 residues |
| Chemical Composition | $C_{2482}H_{3803}N_{649}O_{635}S_{12}$ |
| Total Atom Count | 7,581 atoms |
| Estimated Biological Half-Life | 1.1 hours (mammalian reticulocytes, *in vitro*) >10 hours (*E. coli*, *in vivo*) |
| Stability Indicator (Instability Index) | 26.51 – classified as stable |
| Hydropathicity Score (GRAVY) | 0.365 |

**Table 12.**
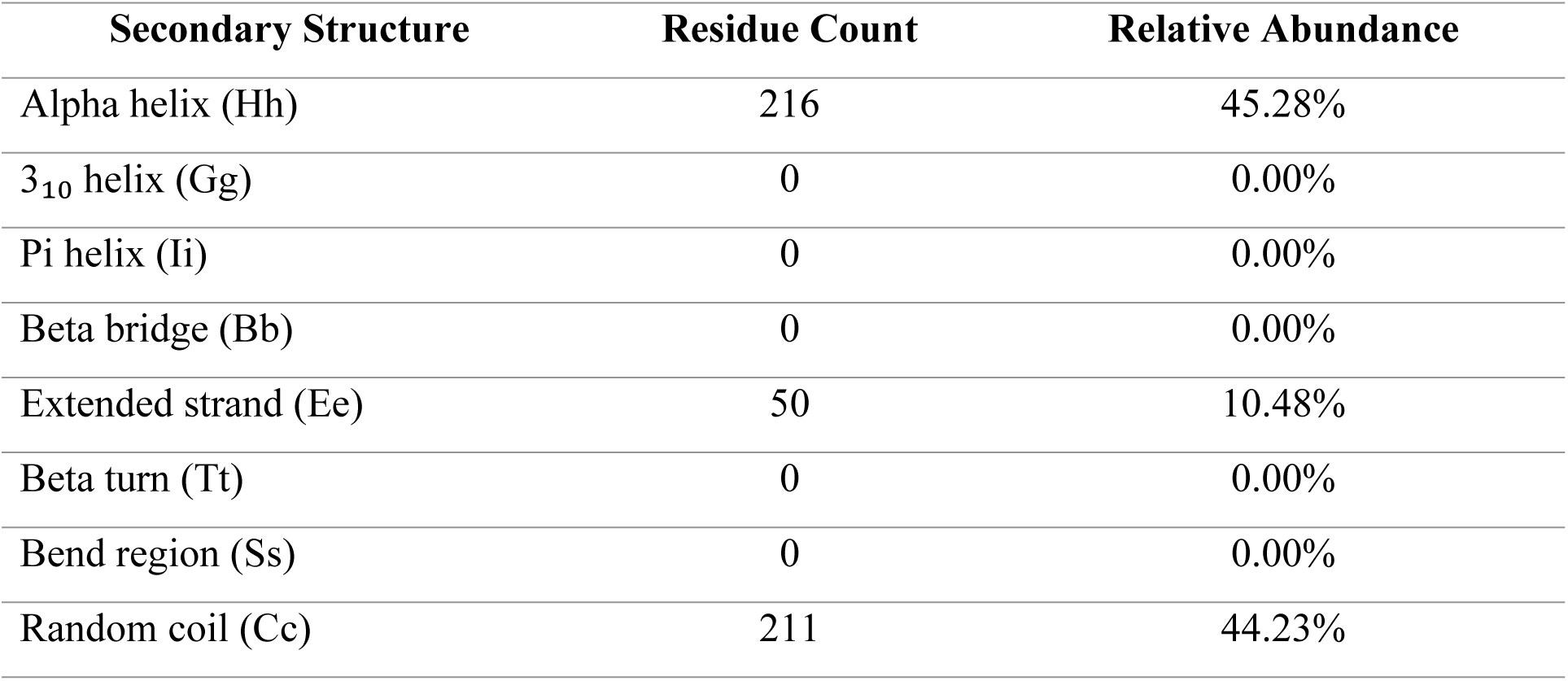
SOPMA results of vaccine construct.

| Secondary Structure | Residue Count | Relative Abundance |
| --- | --- | --- |
| Alpha helix (Hh) | 216 | 45.28% |
| 3<sub>10</sub> helix (Gg) | 0 | 0.00% |
| Pi helix (Ii) | 0 | 0.00% |
| Beta bridge (Bb) | 0 | 0.00% |
| Extended strand (Ee) | 50 | 10.48% |
| Beta turn (Tt) | 0 | 0.00% |
| Bend region (Ss) | 0 | 0.00% |
| Random coil (Cc) | 211 | 44.23% |

**Table 13.** Parameters such as GDT-HA, RMSD, MolProbity score, clash score, rotamer quality, and Ramachandran plot analysis of refined models.

| Model | GDT-HA | RMSD | MolProbity | Clash score | Poor rotamers | Rama favored |
| --- | --- | --- | --- | --- | --- | --- |
| Initial | 1 | 0 | 3.578 | 87.6 | 8.7 | 92.8 |
| MODEL 1 | 0.8863 | 0.565 | 1.683 | 11.6 | 0.3 | 97.5 |
| MODEL 2 | 0.8842 | 0.579 | 1.693 | 10.9 | 0.5 | 97.3 |
| MODEL 3 | 0.8884 | 0.564 | 1.698 | 10.3 | 0.3 | 97.1 |
| MODEL 4 | 0.8747 | 0.582 | 1.669 | 10.3 | 0.5 | 97.3 |
| MODEL 5 | 0.8884 | 0.574 | 1.629 | 9.2 | 0 | 97.3 |

**Table 14.**
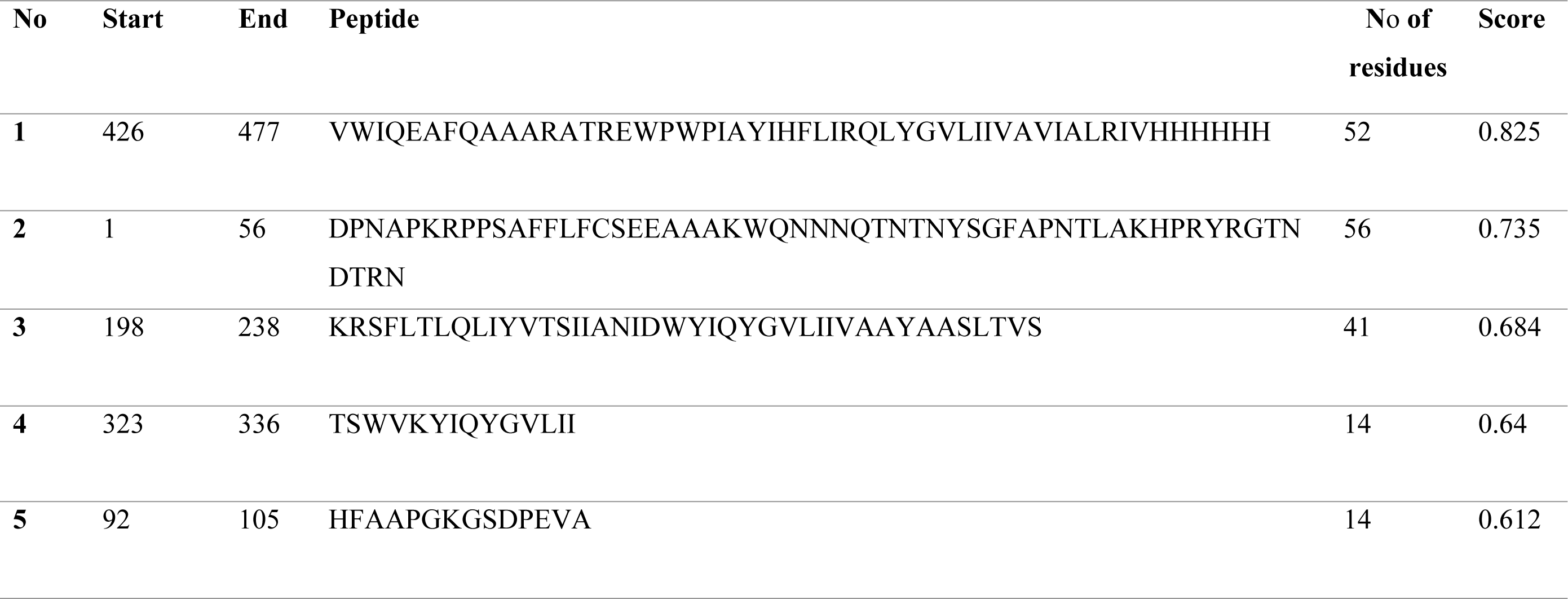
Five high-scoring linear epitopes were identified.

**Table 15.** Protein–Protein docking results of vaccine construct and receptors 3FXL.

| Cluster | Members | Representative | Weighted Score |
| --- | --- | --- | --- |
| 1 | 62 | Center | -1374.3 |
|  |  | Lowest Energy | -1572.2 |
| 2 | 30 | Center | -1086.8 |
|  |  | Lowest Energy | -1378.9 |
| 3 | 29 | Center | -1396.7 |
|  |  | Lowest Energy | -1396.7 |
| 4 | 26 | Center | -1131.7 |
|  |  | Lowest Energy | -1364.8 |
| 5 | 26 | Center | -1302 |
|  |  | Lowest Energy | -1439.8 |
| 6 | 25 | Center | -1409.8 |
|  |  | Lowest Energy | -1409.8 |

**Table 16.**
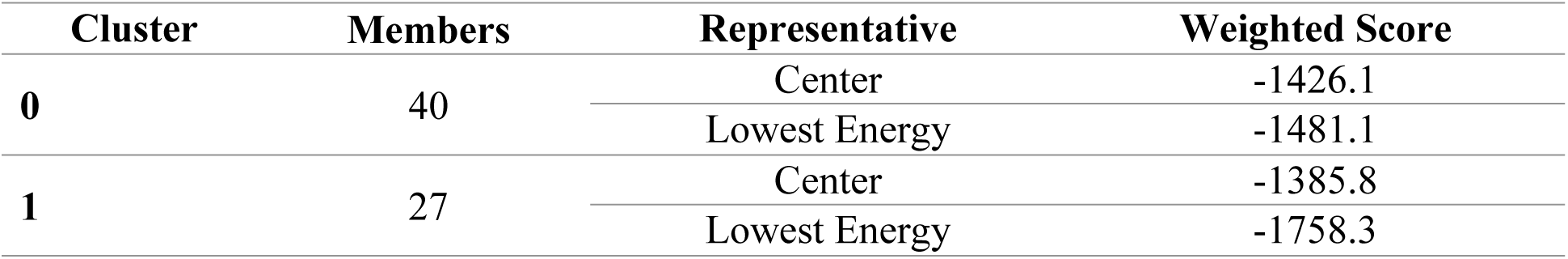
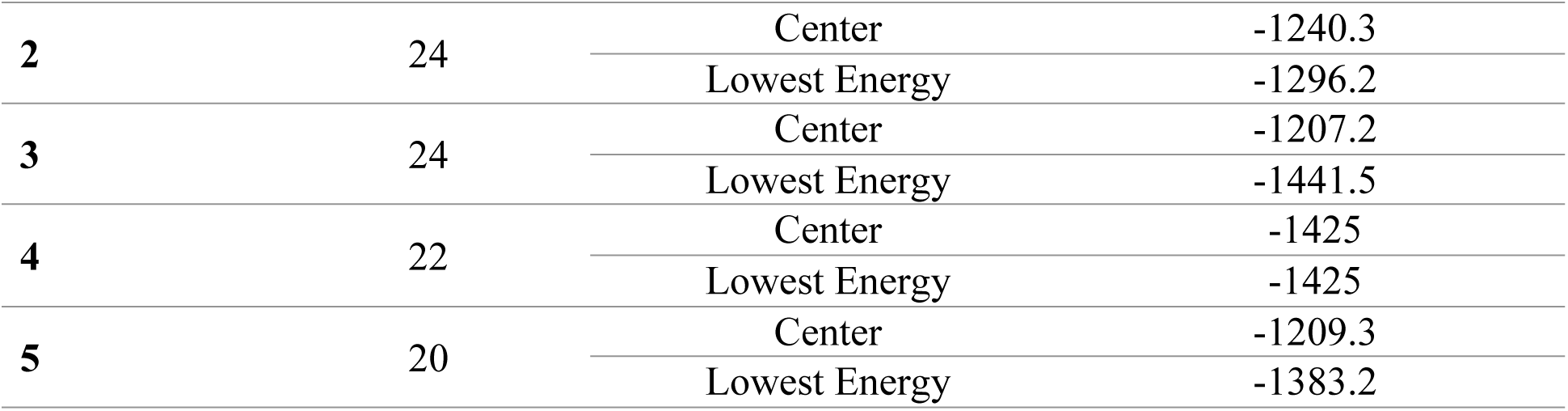
Protein docking results of vaccine construct and receptors 2AOZ.

**Table 17.**
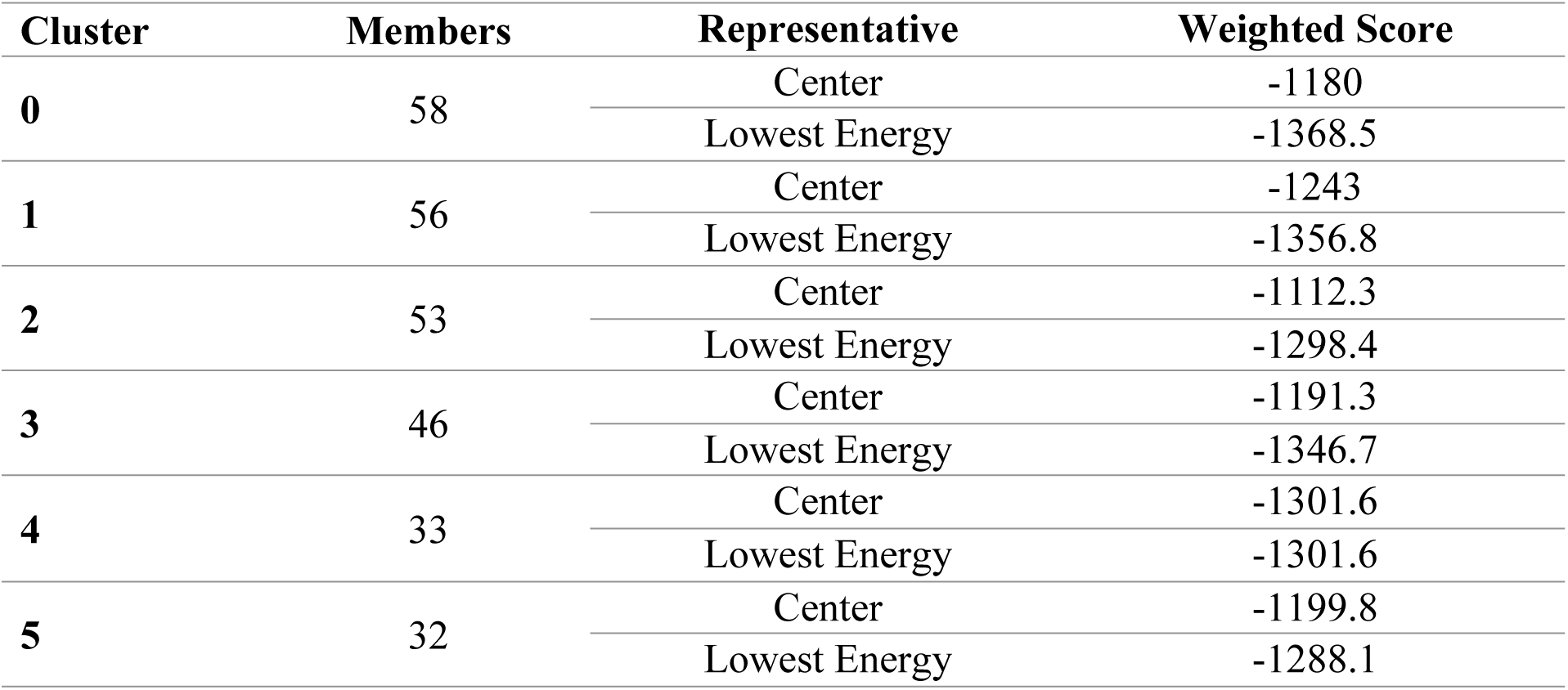
Protein–Protein docking results of vaccine construct and receptors 1Z1W.

| **Cluster** | **Members** | **Representative** | **Weighted Score** |
| --- | --- | --- | --- |
| **0** | 58 | Center | -1180 |
|  |  | Lowest Energy | -1368.5 |
| **1** | 56 | Center | -1243 |
|  |  | Lowest Energy | -1356.8 |
| **2** | 53 | Center | -1112.3 |
|  |  | Lowest Energy | -1298.4 |
| **3** | 46 | Center | -1191.3 |
|  |  | Lowest Energy | -1346.7 |
| **4** | 33 | Center | -1301.6 |
|  |  | Lowest Energy | -1301.6 |
| **5** | 32 | Center | -1199.8 |
|  |  | Lowest Energy | -1288.1 |

**Table 18.** Protein–Protein docking results of vaccine construct and receptors 3WPB.

| **Cluster** | **Members** | **Representative** | **Weighted Score** |
| --- | --- | --- | --- |
| **0** | 26 | Center | -1392.3 |
|  |  | Lowest Energy | -1669.3 |
| **1** | 25 | Center | -1326.7 |
|  |  | Lowest Energy | -1509.7 |
| **2** | 24 | Center | -1421.8 |
|  |  | Lowest Energy | -1558.3 |
| **3** | 21 | Center | -1578.7 |
|  |  | Lowest Energy | -1610 |
| **4** | 20 | Center | -1347.5 |
|  |  | Lowest Energy | -1676.6 |
| **5** | 20 | Center | -1409.1 |
|  |  | Lowest Energy | -1754.9 |

**Table 19.** Summary of MD simulation parameters describing the structural stability and conformational behavior of the protein–protein complex. Parameters include RMSD, RMSF, Rg, and SASA, with corresponding values and interpretations over the simulation timeline.

| Parameter | Meaning | Value/Range | Interpretation |
| --- | --- | --- | --- |
| RMSD | Assesses the structural deviation from the initial conformation | Minor variation around 40 ns; remains steady afterward | Reflects the structural stability of complex (Figure 14a) |
| RMSF | Evaluates residue-level movement and flexibility | Below 3.35 nm, higher at binding regions | Highlights flexible regions and interaction hotspots (Figure 14b) |
| Rg | Measures structural compactness and folding consistency | Maintains stability from 40 ns to 100 ns | Indicates a well-folded, stable conformation (Figure 14c) |
| SASA | Calculates solvent-exposed surface area of the protein | Shows a downward trend until ~20 ns, then plateaus | Suggests the protein surface becomes stable during the simulation |

### Epitope Conservation analysis

Conservancy analysis using IEDB showed that most epitopes matched only 1 out of 13 full-length protein sequences at 100% identity, indicating a low conservation rate (7.69%). MHC-I epitopes such as AMGAASLTV had minimal identity ranging from 10–22.22%. MHC-II epitopes like PIAYIHFLIRQLIRL showed slightly better conservation (∼20%) but were still limited to one sequence match (Table 7 and 8).

### Population coverage analysis

The frequency of HLA alleles varies significantly across diverse ethnic groups. The top scoring epitopes were selected for population coverage percentages were indicated (Tables 2 and 3) against allele of MCH-I and MCH-II. The population coverage was of west Africa, Pakistan and global are showing. The population coverage of West Africa of MCH I is 69.51%, Pakistan is 78%% and global is 82%. The population coverage of West Africa of MCH II is 99%, Pakistan is 1% and global is 99%.

### Design of multiepitope peptide constructs

The vaccine construct was made with adjuvant of HP91 with sequence DPNAPKRPPSAFFLFCSE, linker EAAAK then connect with B cell epitopes. A linker of CPGPG connect the MHC-I epitopes with B cell epitopes and linker of AAYA is present between MCH-II epitopes and MCH-II epitopes end with 6X Histadine Tag HHHHHH.The Overall Prediction for the Protective Antigen = 0.7146 (Probable ANTIGEN) and Threshold for this model was 0.45 and Allertop v2.1 showed Probable NON-ALLERGEN. The whole sequence is shown below

### Physicochemical and Immunological Profile of Vaccine Constructs

The results obtained from the ProtParam server are presented in Table 4. The designed vaccine constructs exhibited an average molecular weight of approximately 53 kDa, which falls within the acceptable range for subunit vaccines. All constructs were predicted to be soluble in the human host, enhancing their potential for efficient expression and delivery. Furthermore, immunoinformatics analysis confirmed that each construct was antigenic and non-allergenic, supporting their suitability for further experimental validation.

### Secondary structure

The secondary structure of the multi-epitope vaccine construct was analyzed using SOPMA and PSIPRED servers. SOPMA results (Table 5) showed that alpha-helices comprise approximately 45% of the structure, making them the dominant secondary element. Beta-sheets and random coils were also present, distributed throughout the sequence as illustrated in Figure 3.

**Figure 1.**
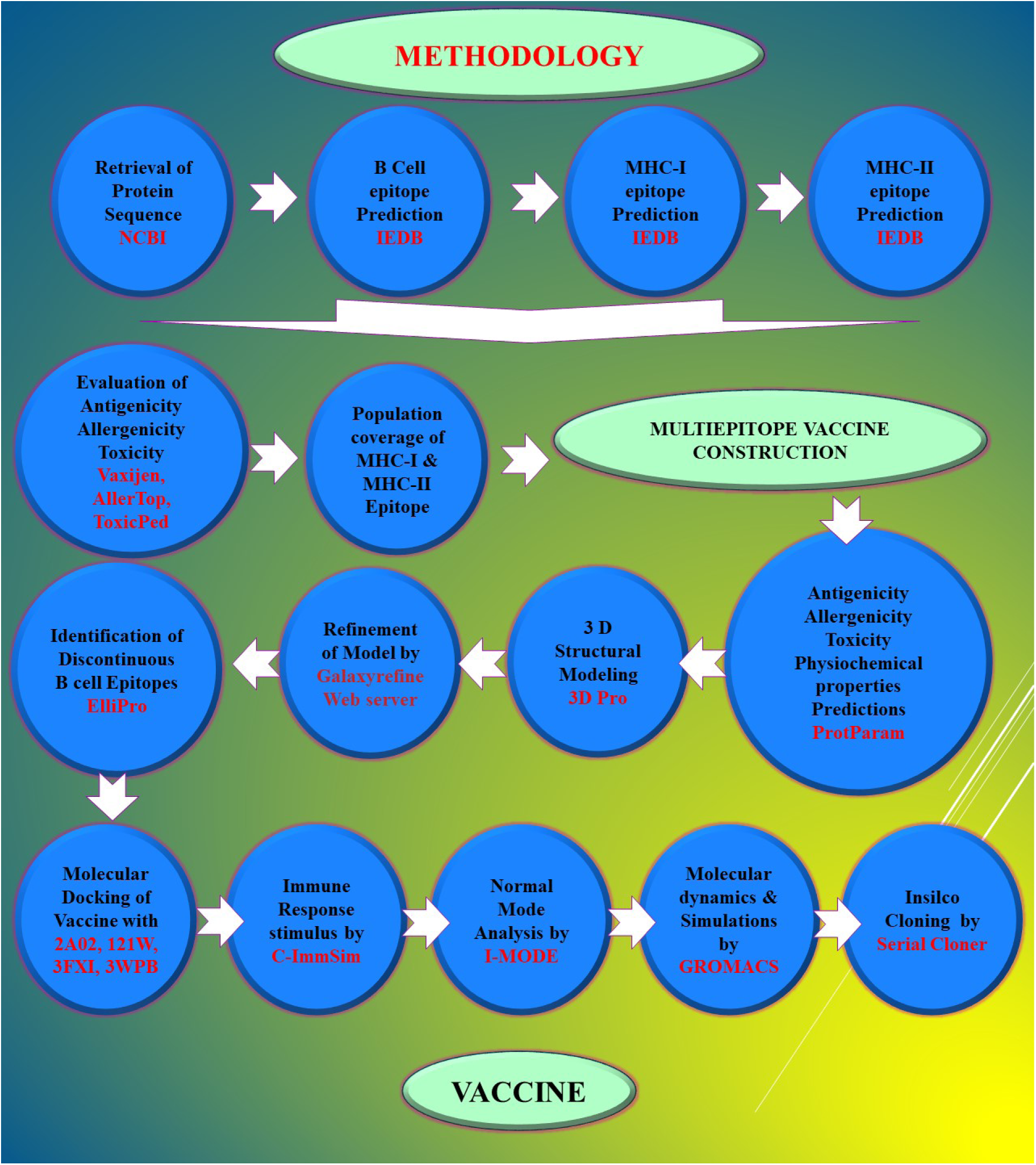
Stepwise complete methodology used in this study

**Figure 2.**
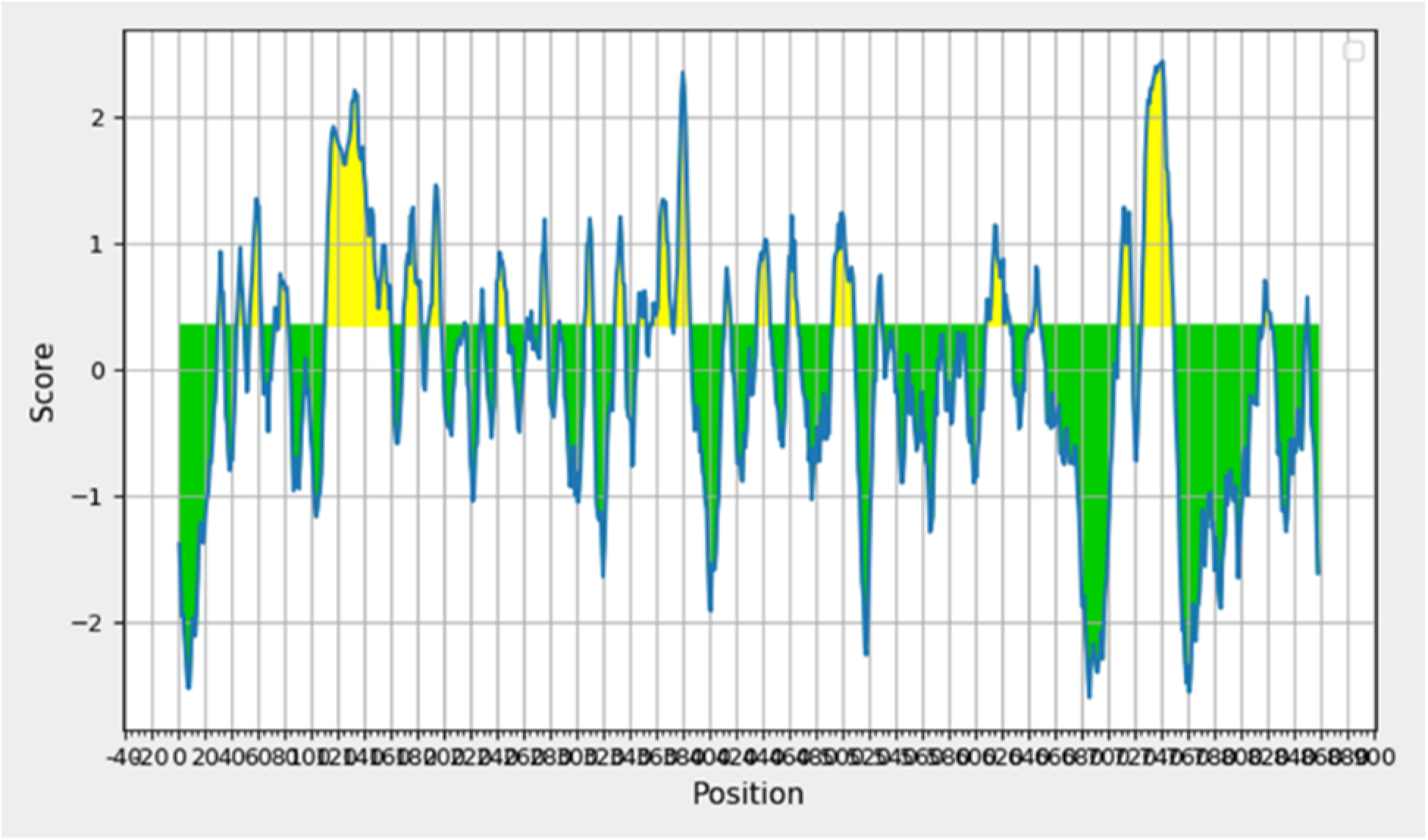
Graph showing the B-cell epitopes predicted for the HIV-2 nev proteins.

**Figure 3.**
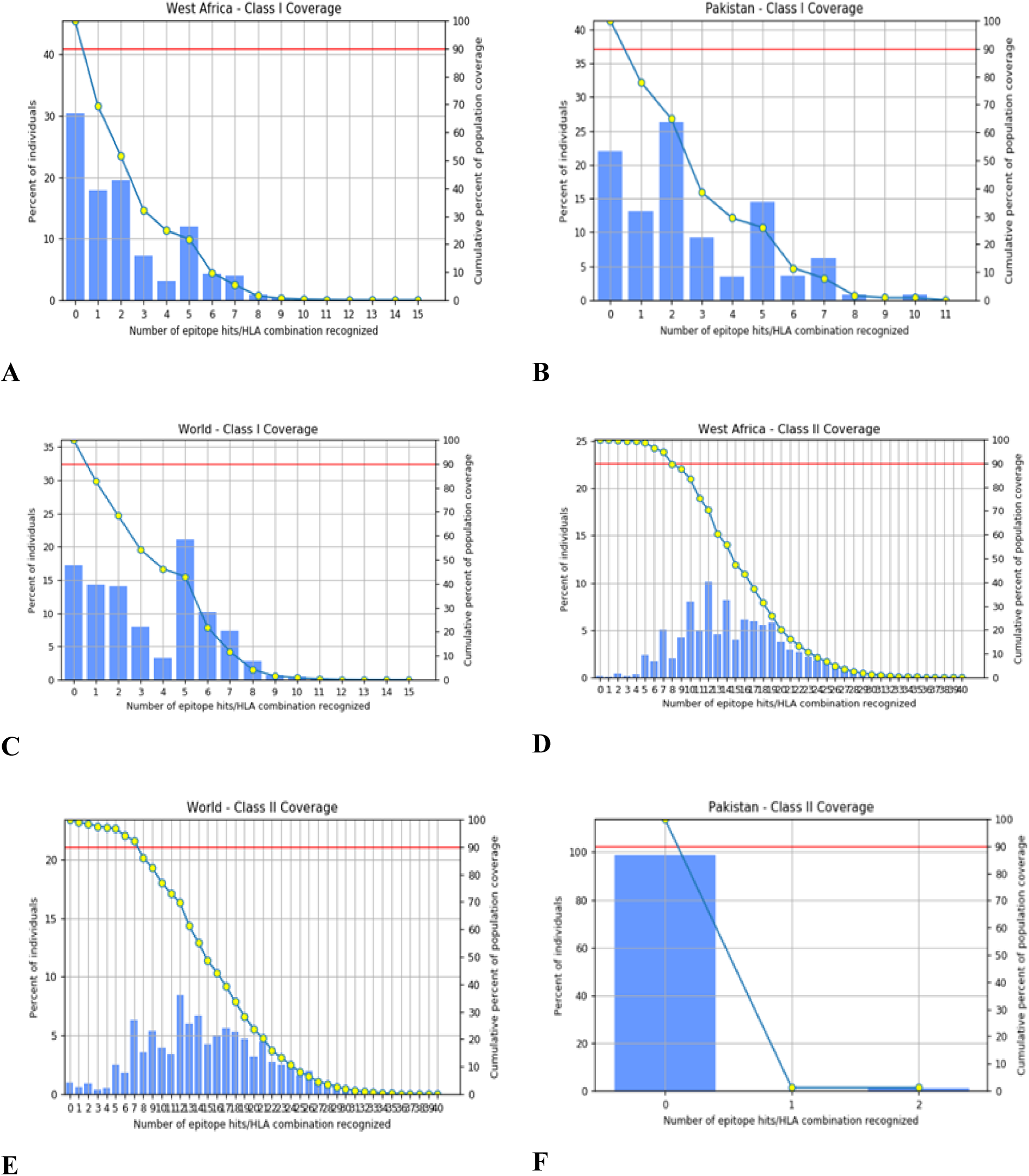
Population coverage analysis: A) The population coverage of West Africa of MCH I is 69.51%, B) Pakistan is 78%% C) Global is 82%, D) The population coverage of West Africa of MCH II is 99%, E) Pakistan is 1% F) Global is 99%.

This balanced arrangement of alpha-helices, beta-sheets, and coil regions suggests proper folding and structural stability. The coil regions may enhance the surface exposure of epitopes, which is crucial for effective immune recognition. Overall, the secondary structure profile indicates that the vaccine construct possesses the necessary structural integrity and flexibility to elicit a strong immunological response.

The results of psipred with A detailed secondary structure annotation of the vaccine construct (Figure 7) was performed to assess its structural conformation. The prediction revealed a predominance of alpha-helical structures (pink), interspersed with random coils (white) and a few beta-sheet elements (blue). Key functional residues, including cysteines (yellow) potentially involved in disulfide bridge formation, were also highlighted. The flexible coil regions are Likely to enhance surface exposure of epitopes, contributing to immunogenicity. This structural organization supports the stability and accessibility required for an effective multi-epitope vaccine.

**Figure 4.**
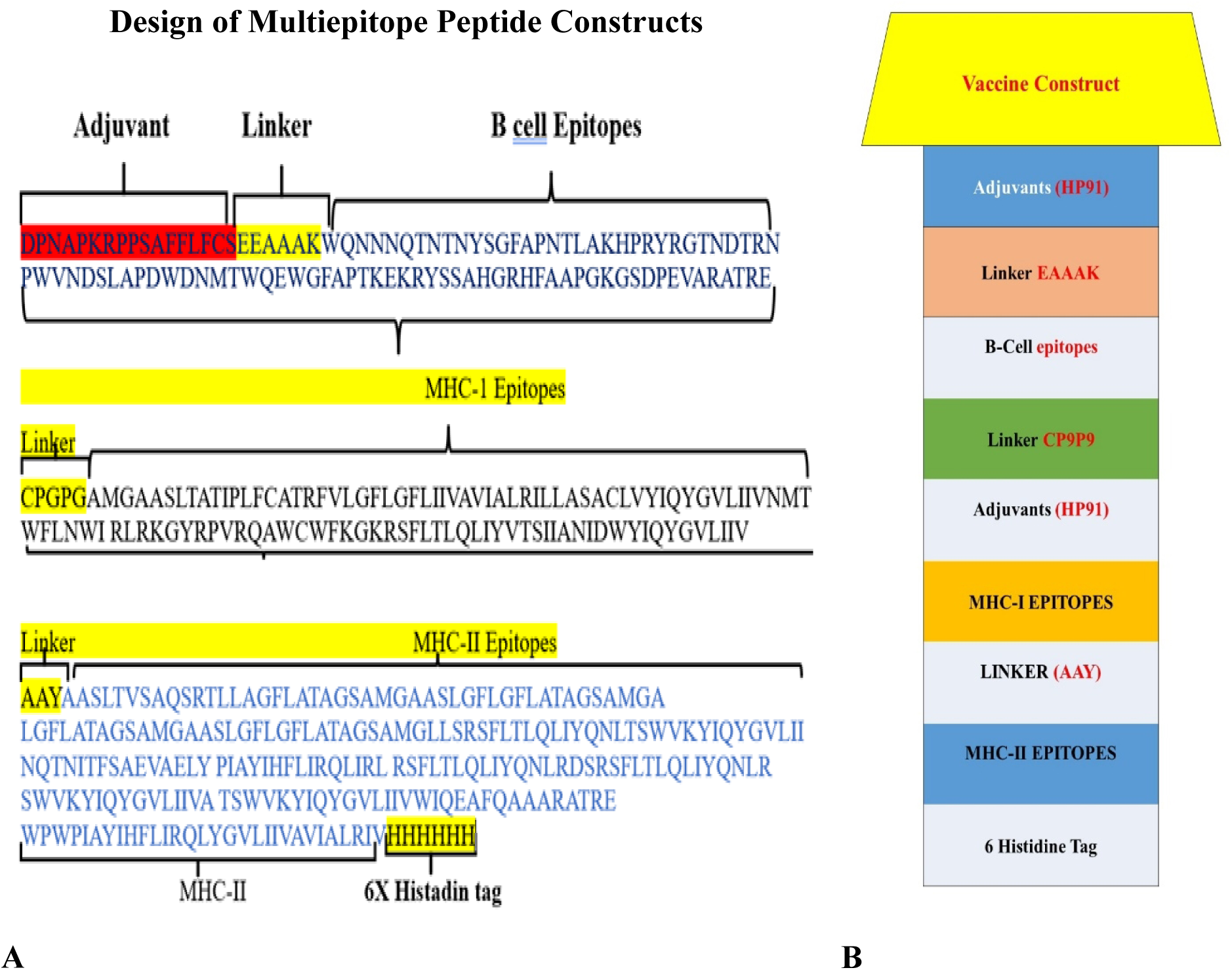
Construct of the vaccines (A) multiple epitope vaccine (B) Linking process

**Figure 5.**
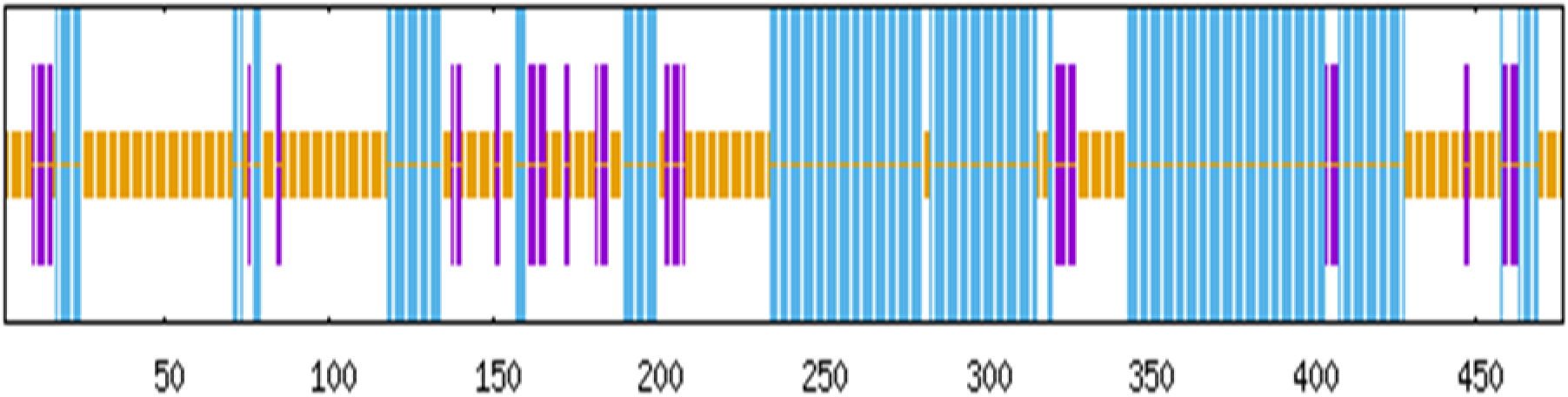
The bar diagram illustrates the predicted secondary structural distribution of the vaccine construct.

**Figure 6.**
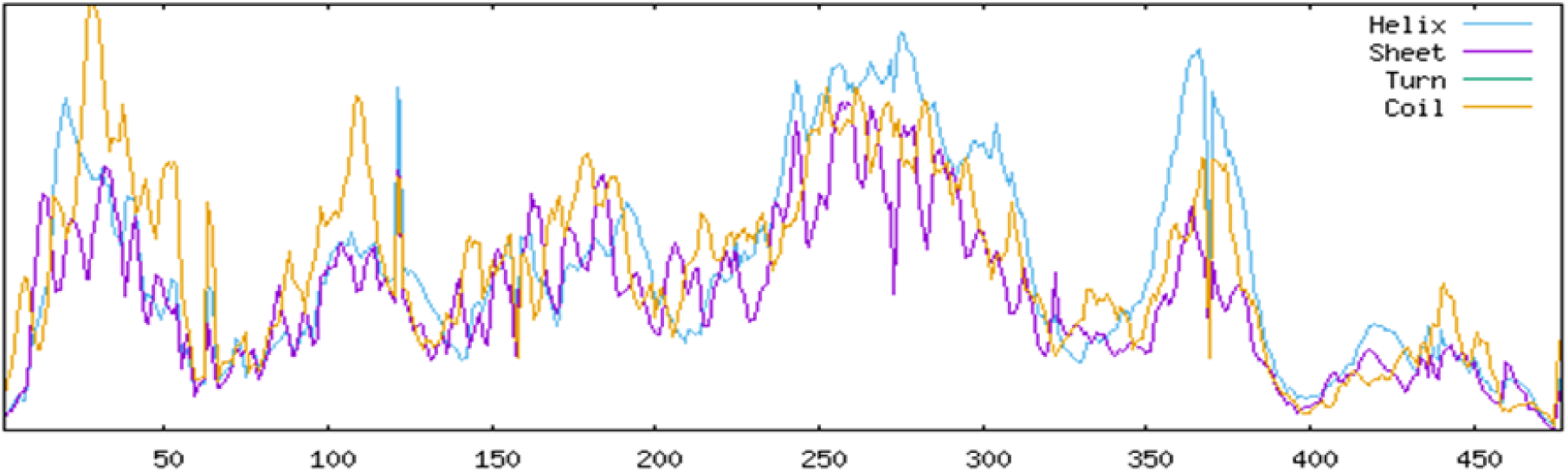
Construction consists of α-helices (light blue), β-sheets (purple), turns (magenta), and random coils (orange) distributed throughout the amino acid sequence.

**Figure 7.**
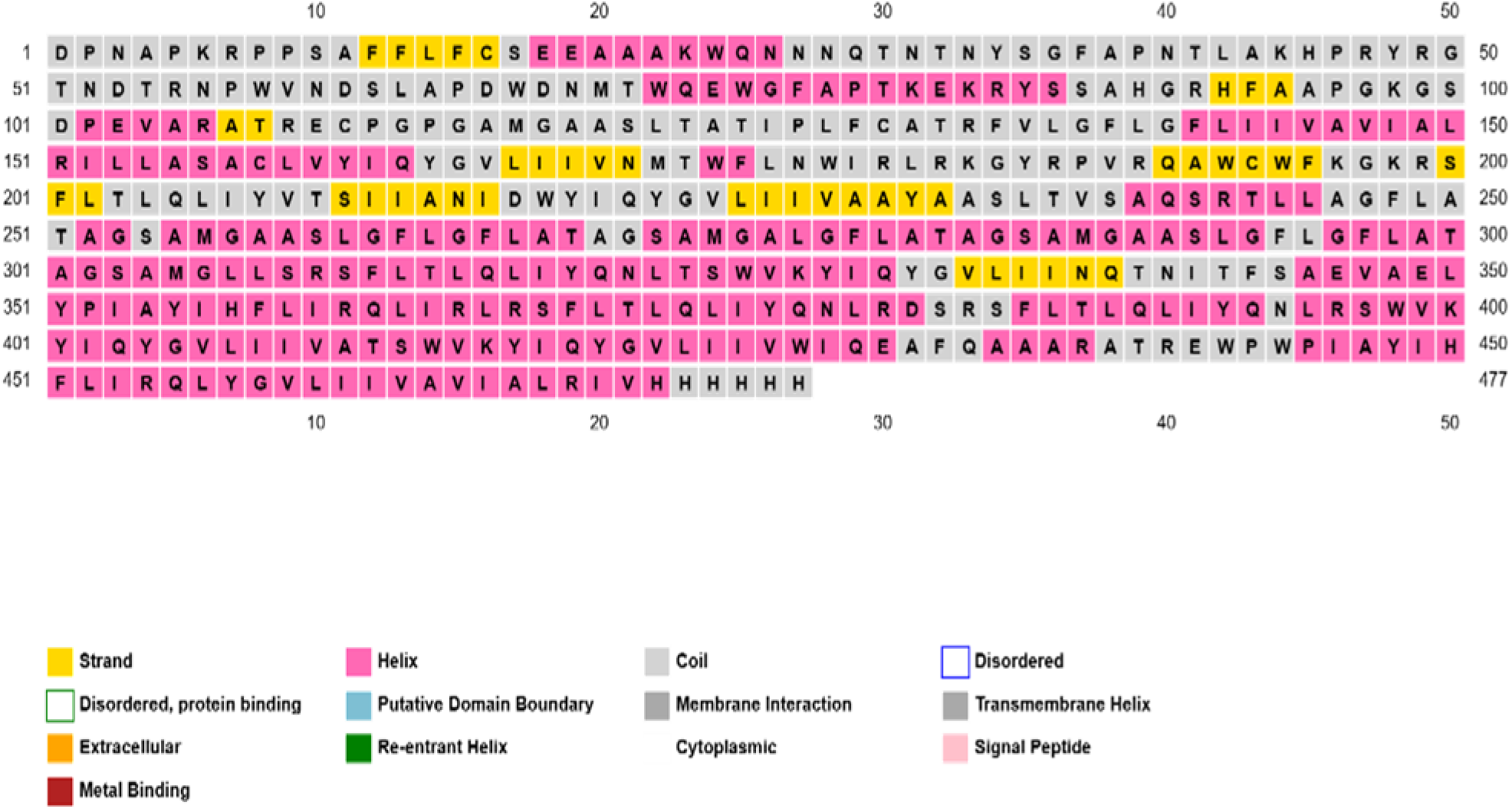
structural conformation

### Predicted tertiary structure

The tertiary structure of the designed multi-epitope vaccine was predicted using the 3Dpro tool available in the SCRATCH protein prediction suite. The predicted model (Figure 8) highlights the overall folding of the construct, dominated by alpha-helices and loop regions. This structure was generated in the absence of suitable templates, utilizing ab initio modeling approaches. The resulting model provides a basis for further structural validation, molecular docking, and dynamic simulation analyses to assess the stability and epitope exposure of the vaccine candidate.”

**Figure 8.**
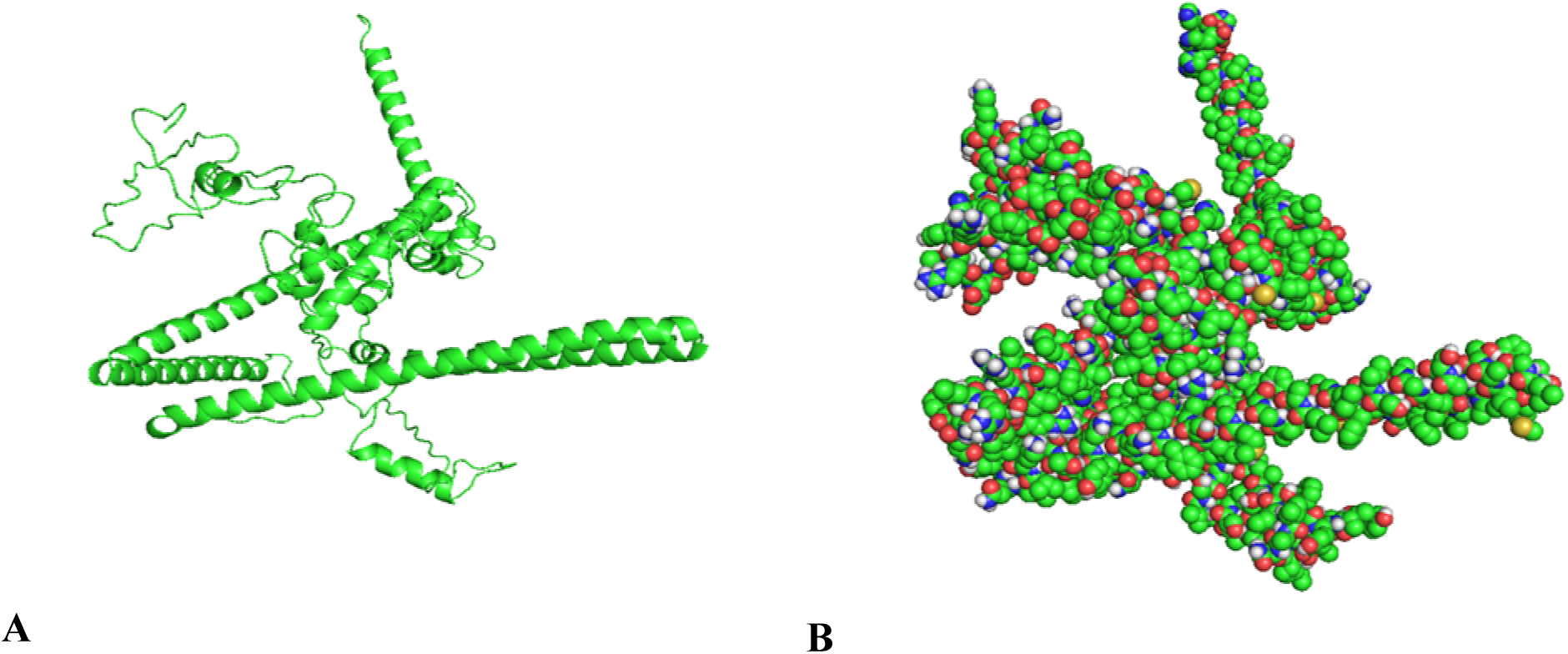
The predicted model highlights the overall folding of the construct, dominated by alpha-helices and loop regions, A is cartoon model and B is surface model.

### Validated 3D structure

The predicted tertiary structure of the vaccine construct was refined and validated using standard quality assessment metrics. Five refined models (Table 6) were evaluated and compared to the initial unrefined model. Parameters such as GDT-HA, RMSD, MolProbity score, clash score, rotamer quality, and Ramachandran plot analysis were used to assess structural integrity and stereochemical quality.The initial model displayed a high MolProbity score (3.578) and clash score (87.6), indicating poor stereochemical quality. Upon refinement, Model 1 showed the best overall results with Ramachandran favored region percentage (97.5%), MolProbity score (1.683), 0.3 rotamers score and with clash score (11.6). Although the GDT-HA score remained relatively consistent across refined models (∼0.88), improvements in RMSD and stereochemical parameters confirm enhanced structural quality. Hence, Model 1 was selected as the final model for further downstream analyses such as molecular docking and immune simulation.

The PROCHECK used for validation and we selected the model 1 for further proves because of Overall Quality Factor is ERRAT 84.35 and Ramachandran plot was 96.0% score (Figure 10).

**Figure 9.**
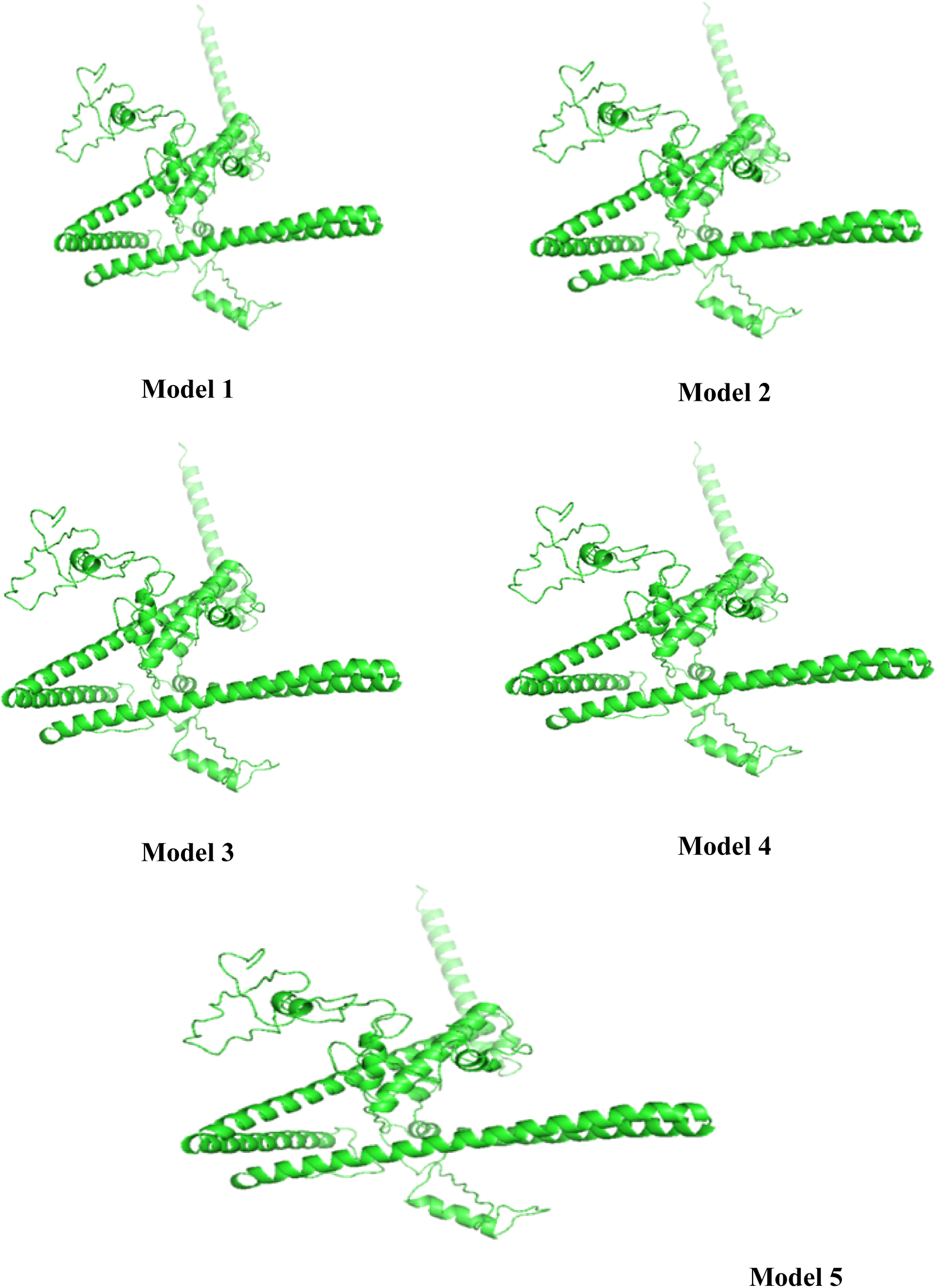
Shows five refined models of vaccine construct

**Figure 10.**
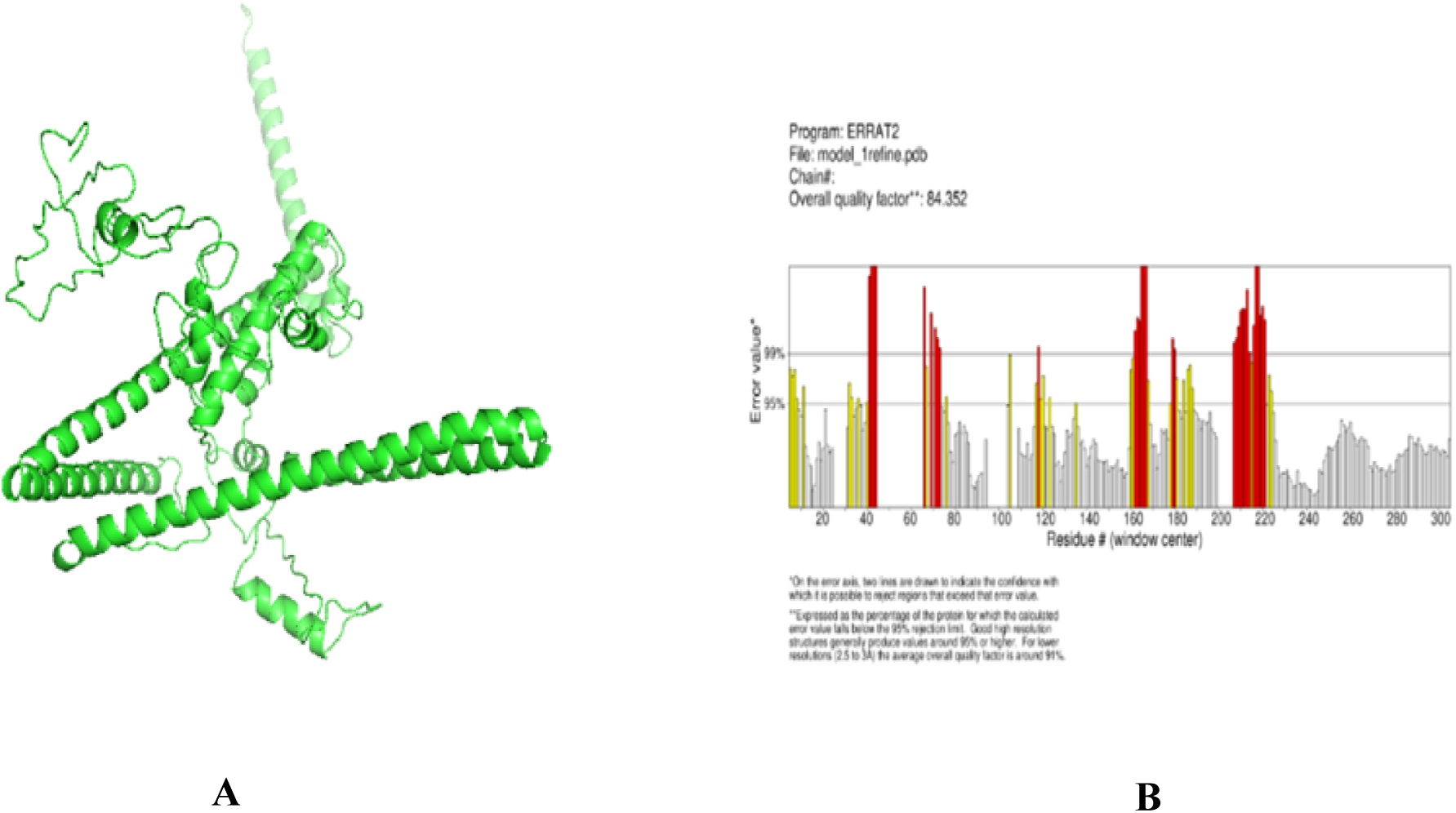
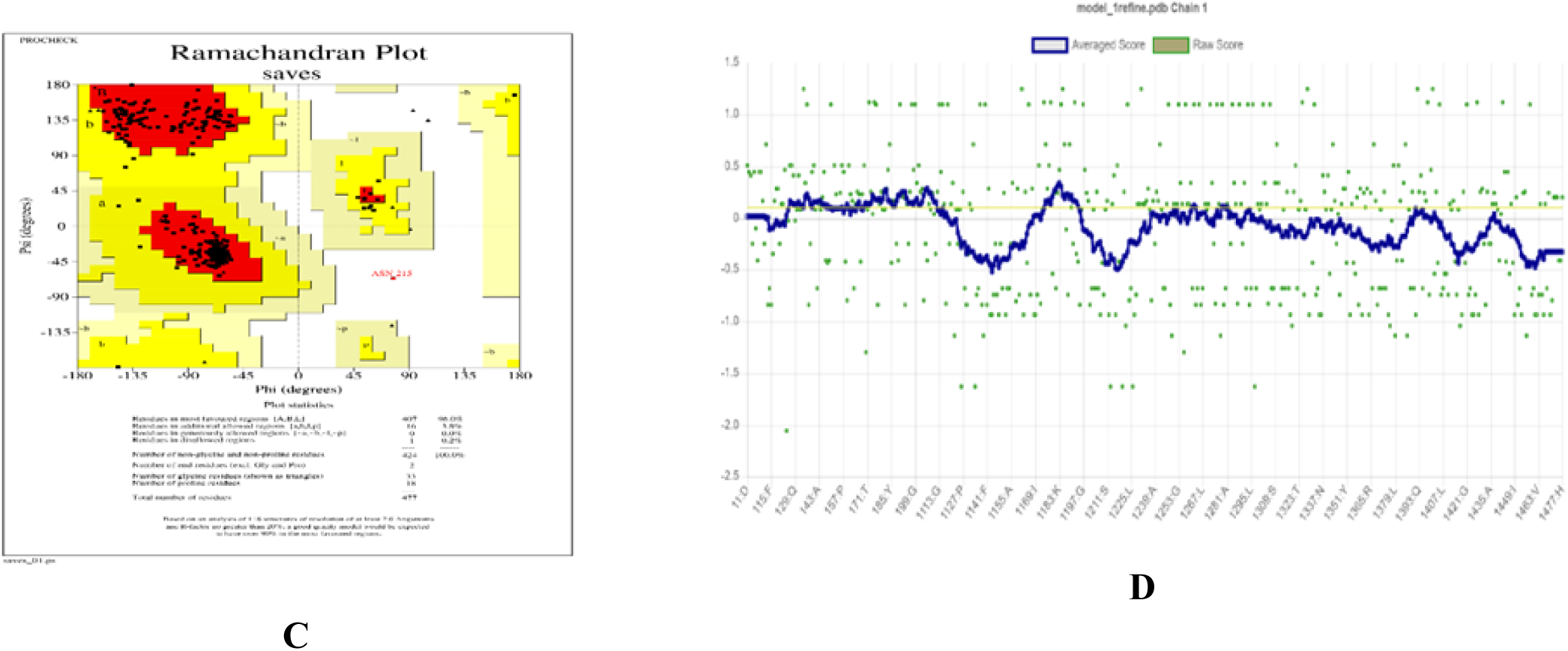
Structural validation of the modeled vaccine construct. The 3D structure was generated using the 3DPro server and refined with the GalaxyRefine server (A). ERRAT analysis yielded an overall quality factor of 84.35% (B). Ramachandran plot analysis using PROCHECK showed that 96.0% of residues were located in the most favored regions (C). VERIFY3D results indicated that 20.34% of the residues had an average 3D-1D score ≥ 0.1 (D).

### Conformational B cell epitope

We Predicted Linear epitope of B-cell which curcial in vaccine design as these regions are recognized by antibodies and contribute to humoral immunity. Five high-scoring linear epitopes were identified (Table 8), ranging from 14 to 56 amino acids in length. The top-ranked epitope (Epitope 1), spanning residues 426 to 477, had the highest prediction score (0.825) and includes a C-terminal His-tag (HHHHHH), which may enhance expression and purification. Epitope 2, found at the N-terminal region (residues 1–56), also showed a high score (0.735), suggesting strong antigenic potential. These predicted epitopes recognized by antibodies, helping the vaccine trigger a strong B-cell-driven immune response.

Linear B-cell epitope prediction conducted by BepiPred 2.0 server also visualized graphically (Figure 11), with scores plotted against the amino acid position of the vaccine construct. The red threshold line (score = 0.5) separates predicted epitopes (above the line) from non-epitopes (below). Several regions in the construct surpassed the threshold, highlighted in yellow, indicating potential antibody-accessible epitopes. Notably, peaks in the C-terminal and N-terminal regions demonstrated high antigenicity, which aligns with previously predicted peptide sequences (see Table 6). The prediction supports the inclusion of these sequences in the final construct for further experimental validation. The graphical visualization confirms the presence of multiple immunodominant regions within the construct, supporting its potential to induce humoral immune responses upon administration.

**Figure 11.**
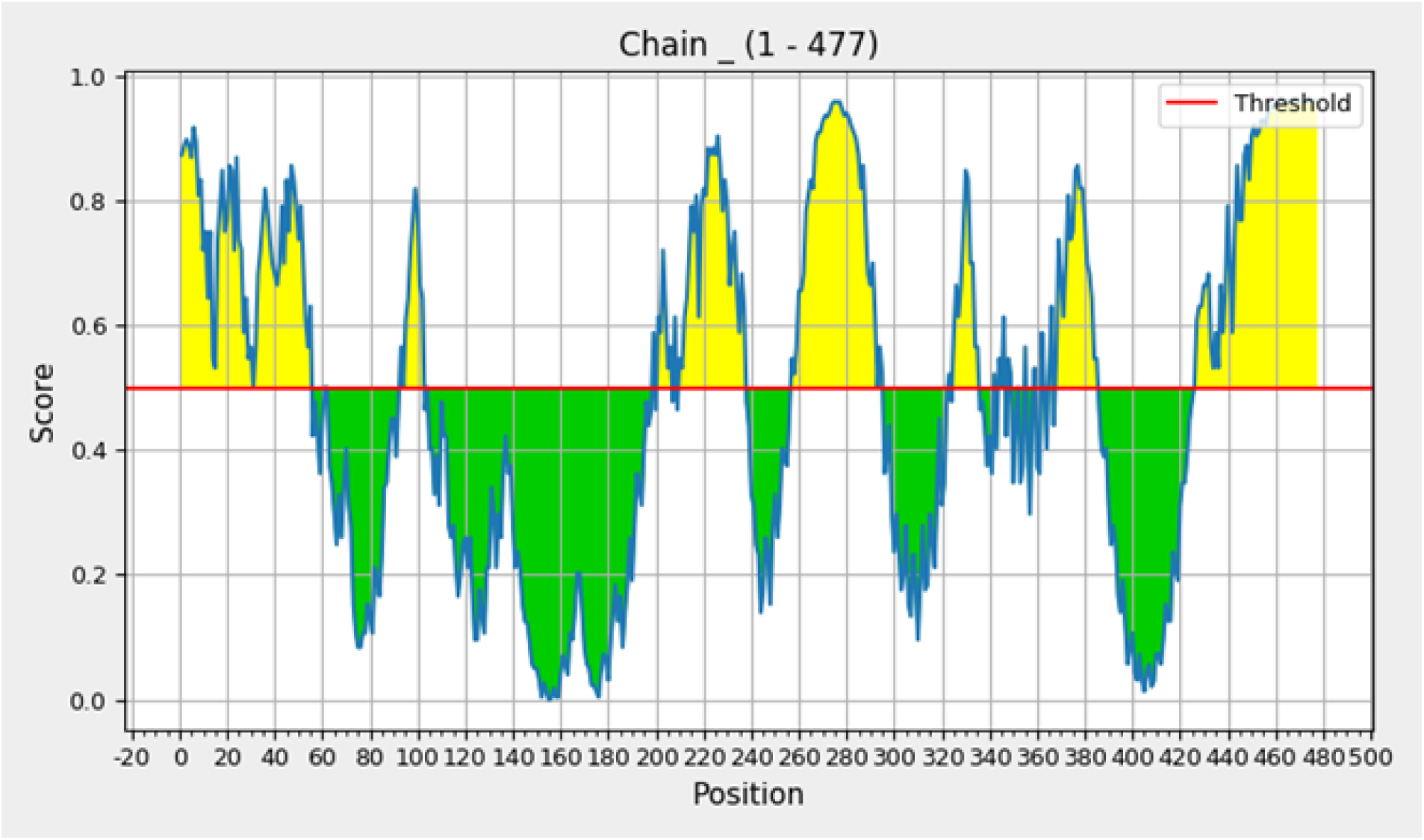
Visualized graphically with scores plotted against the amino acid position of the vaccine construct

### Predicted Discontinuous epitope(s)

Using a default threshold of 0.5, six conformational B-cell epitopes were identified. These epitopes are made up of surface-exposed amino acids that are close together in the 3D structure. Their scores above the prediction cutoff suggest they may effectively trigger antibody responses (Figure X).

### Vaccine Construct and Toll-Like Receptors

The vaccine showed strong binding affinities, with the lowest energy scores of −1481.1 kcal/mol (TLR2), −1572.2 kcal/mol (TLR4), −1368.5 kcal/mol (TLR3), and −1669.3 kcal/mol (TLR9), indicating stable and favorable interactions. Detailed docking analysis with TLR2 (PDB ID: 2AOZ) revealed a robust complex involving Sixteen hydrogen bonds (H-bond), Five salt bridges, and 274 non-bonded contacts. Secondary structure analysis (via PDBsum) indicated a well-defined architecture comprising alpha helices, beta strands, and loops, supporting its structural integrity upon binding. These findings suggest strong molecular compatibility between the vaccine and TLRs, supporting its potential to activate innate immune signaling pathways.

### Structural Flexibility & Normal-Mode Analysis

The deformability plot (Fig. 14A) demonstrated minimal fluctuations, showing a strong structural rigidity. B_factor values obtained from NMA (Figure. 14B) were substantially lower than the experimental B-factors reported in the PDB, further supporting the enhanced stability of the complex. The calculated eigenvalue of 3.93 × 10⁻⁶ indicates that the complex has high molecular flexibility, meaning it can easily undergo structural changes. The covariance matrix (Figure. 14E) shows how different residues move in relation to each other—red indicates they move together, blue means they move oppositely, and white shows no correlation. The elastic network model (Figure. 14F) highlights the strength of atomic connections, where darker gray dots represent stiffer, more tightly connected regions.

**Figure 12.**
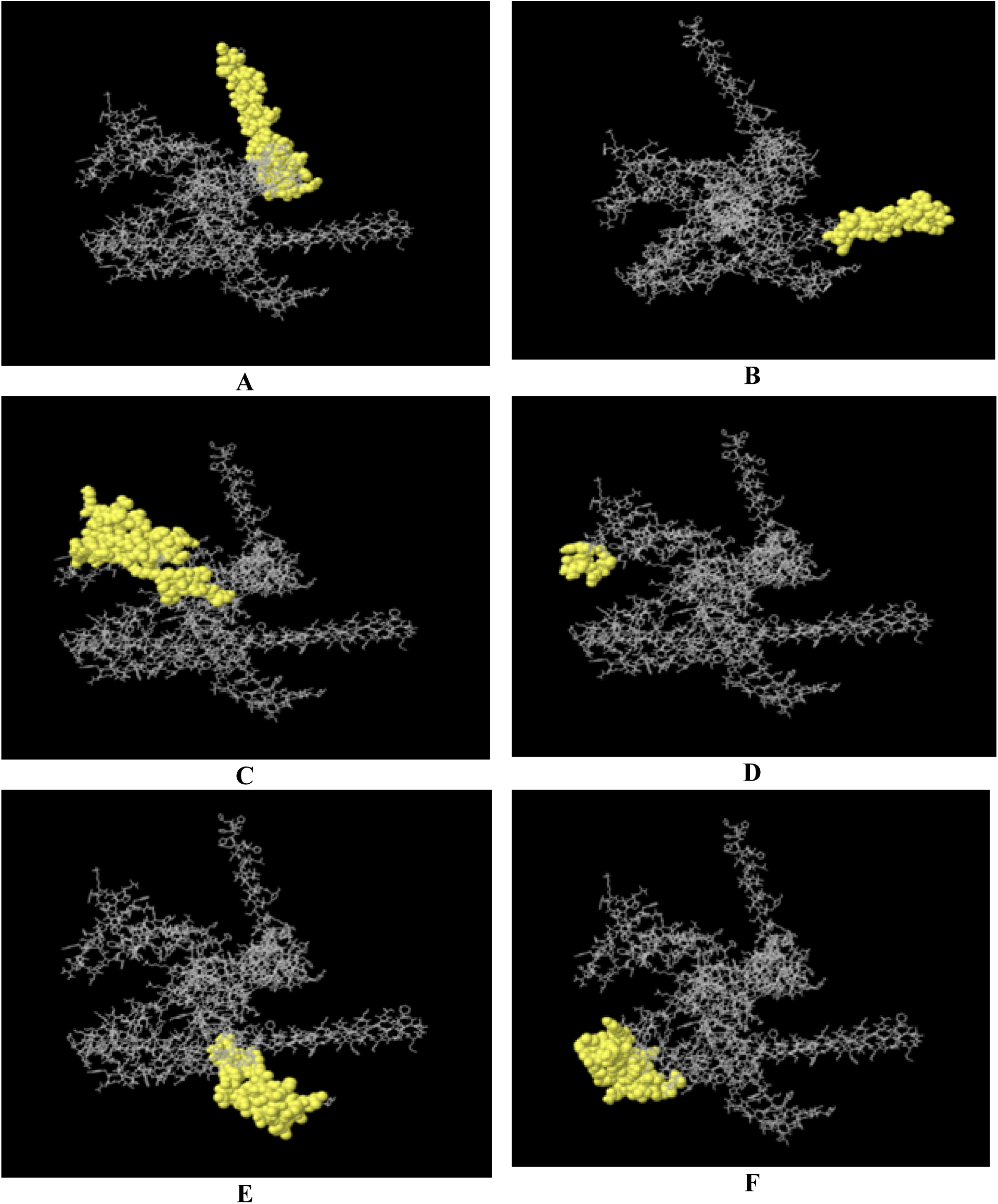
Predicted Discontinuous B-Cell Epitope Mapping: Discontinuous B-cell epitopes mapped onto the vaccine construct (panels a–f) are highlighted in yellow. These epitopes encompass residues 8 to 58, with corresponding score values of 0.882, 0.081, 0.714, 0.708, 0.676, and 0.666, respectively.

**Figure 13.**
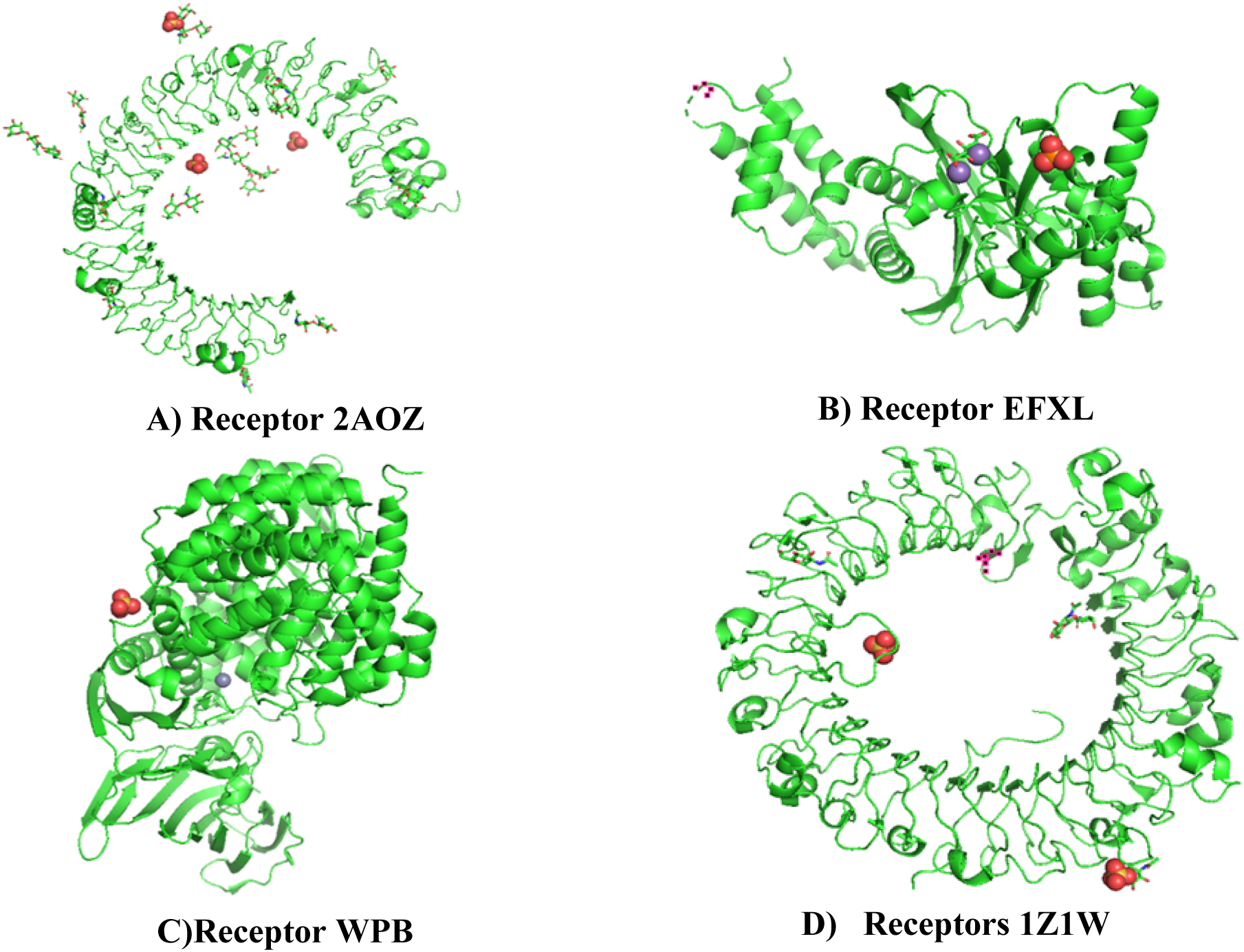
A) TLRs Receptor of 2AOZ, B) 3FXL, C) 31Z1W, D) 3WPB

**Figure 14.**
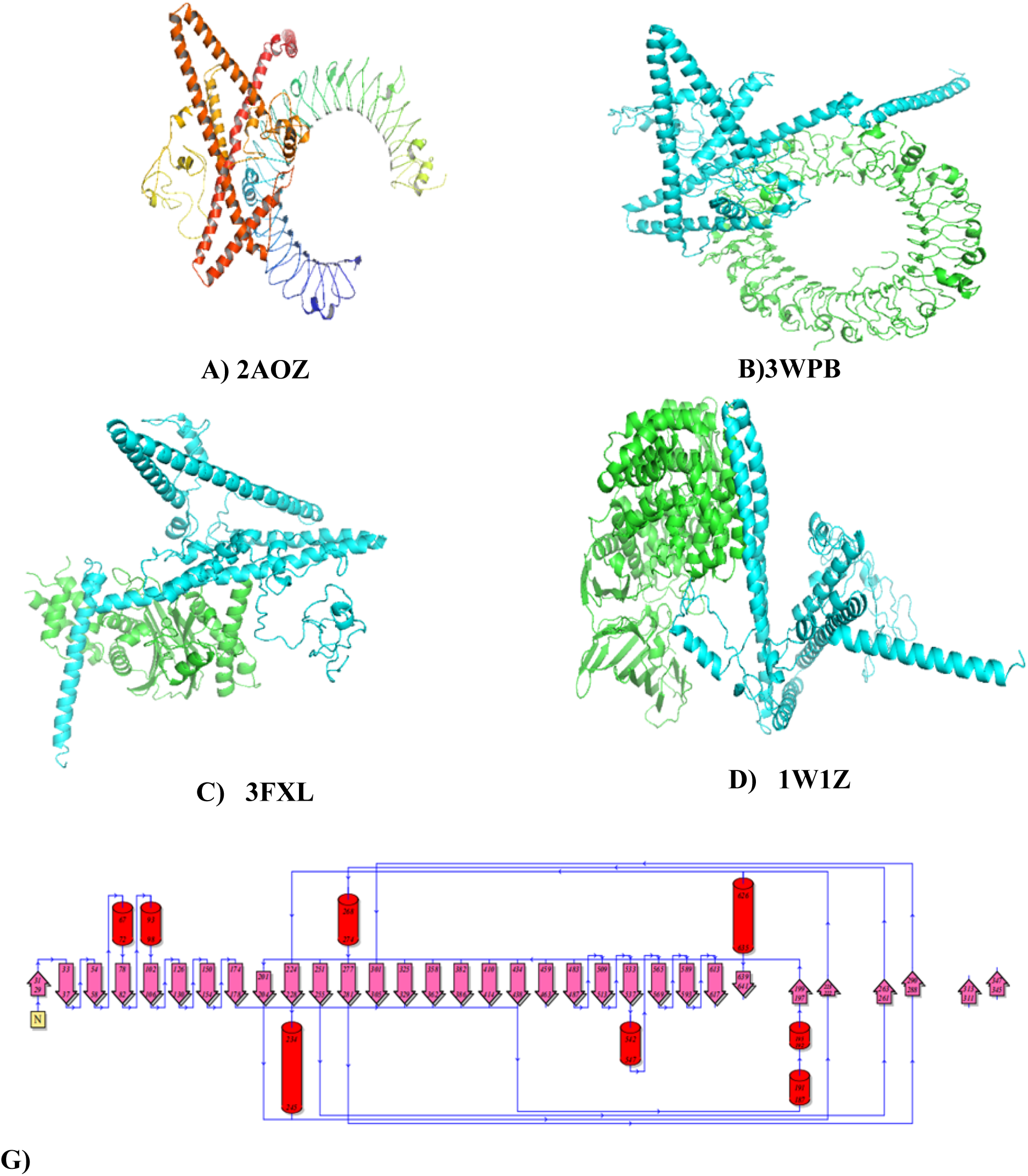
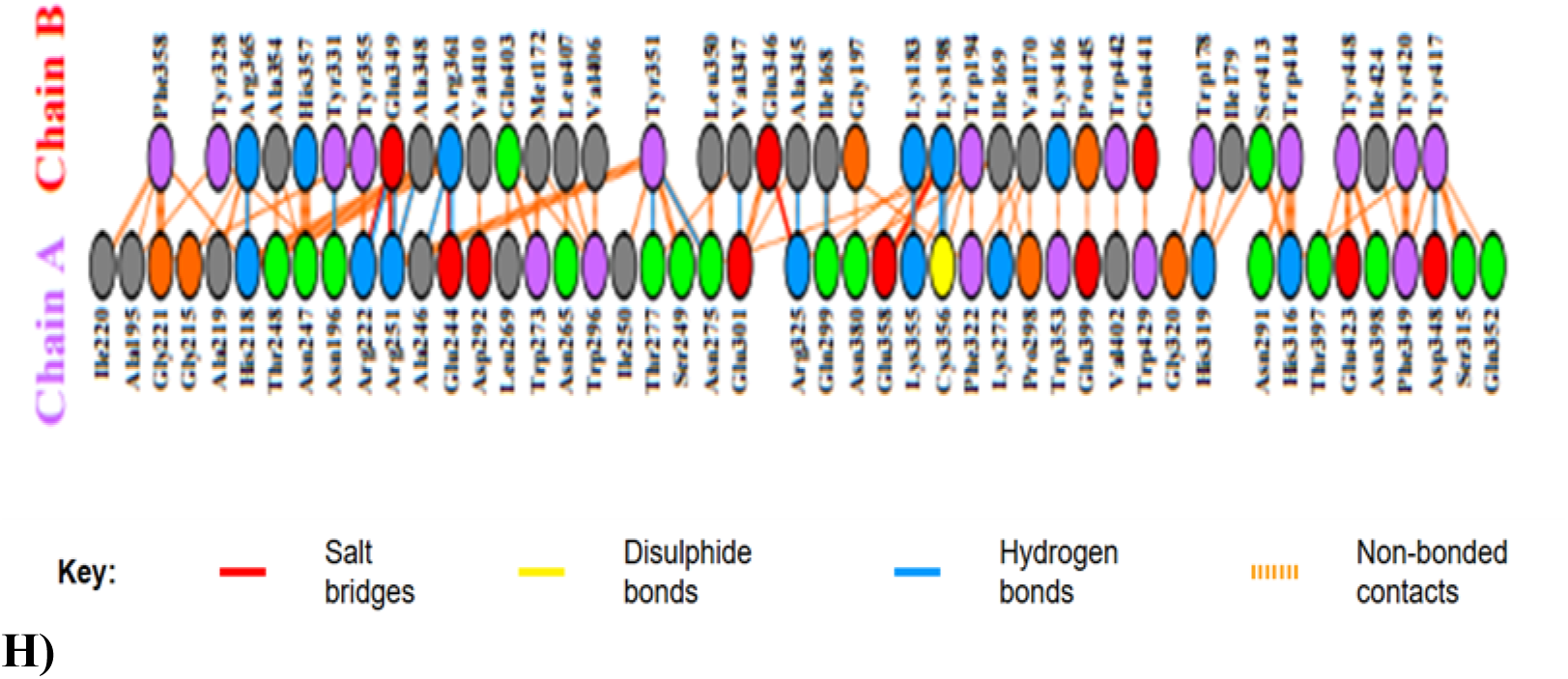
The protein–protein docking between Vaccine construct and TLRs (A; 2AOZ: B; 3WPB: C; 3FXL: D; 1W1Z G) structure of Topology H) binding patterns that are formed in docking complex

**Figure 15.**
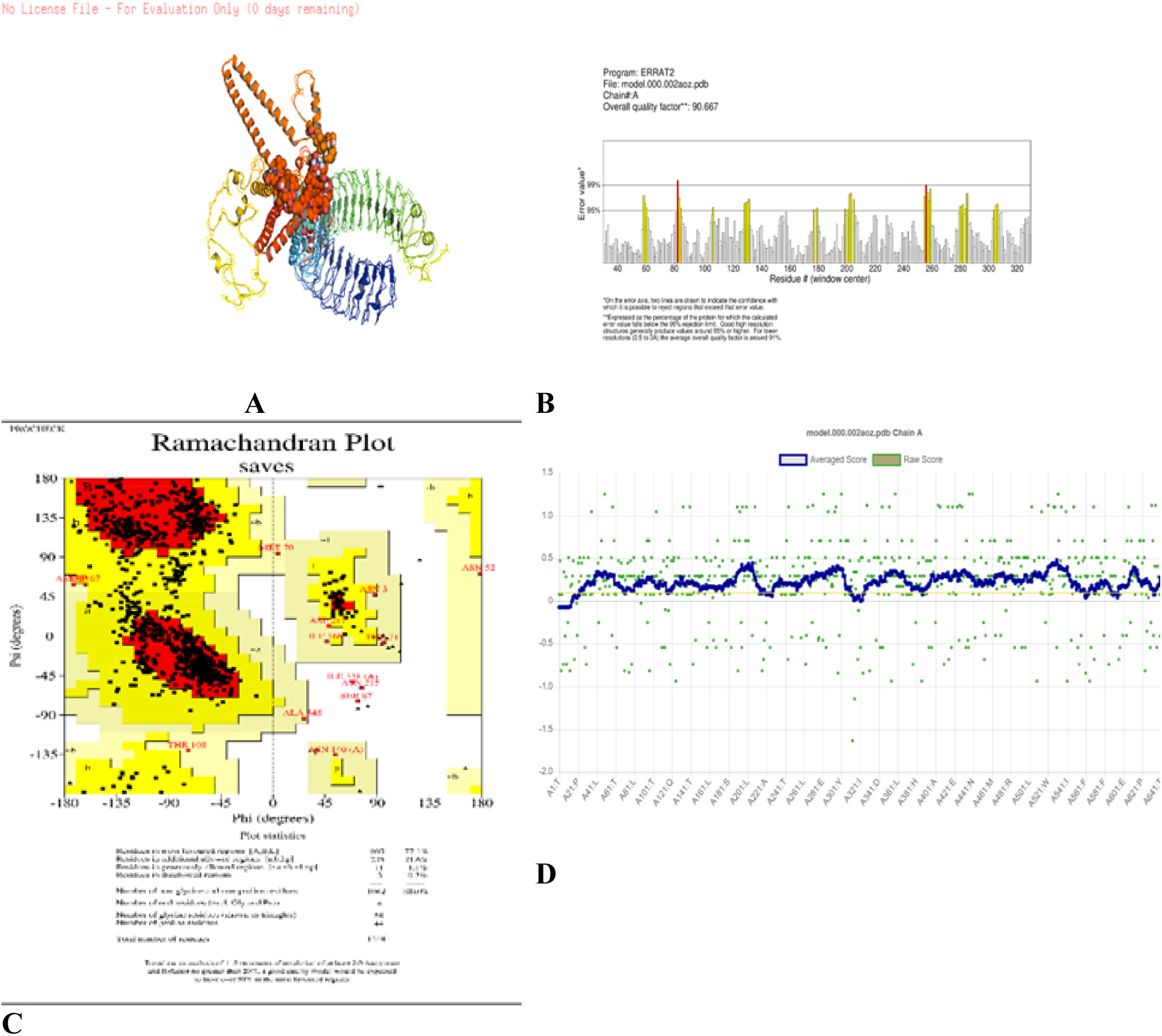
The docking results with Receptor 2AOZ : A) The 3D protein generated by 3DPro Server B) ERRAT error values chart with 90.6667 of quality factor, C) Ramachadran plot analysis by PROCHECK shows 77.1% of residues in the most favoured regions; D) VERIFY3D 91.65% of the residues have averaged 3D-1D score >= 0.1

**Figure 16.**
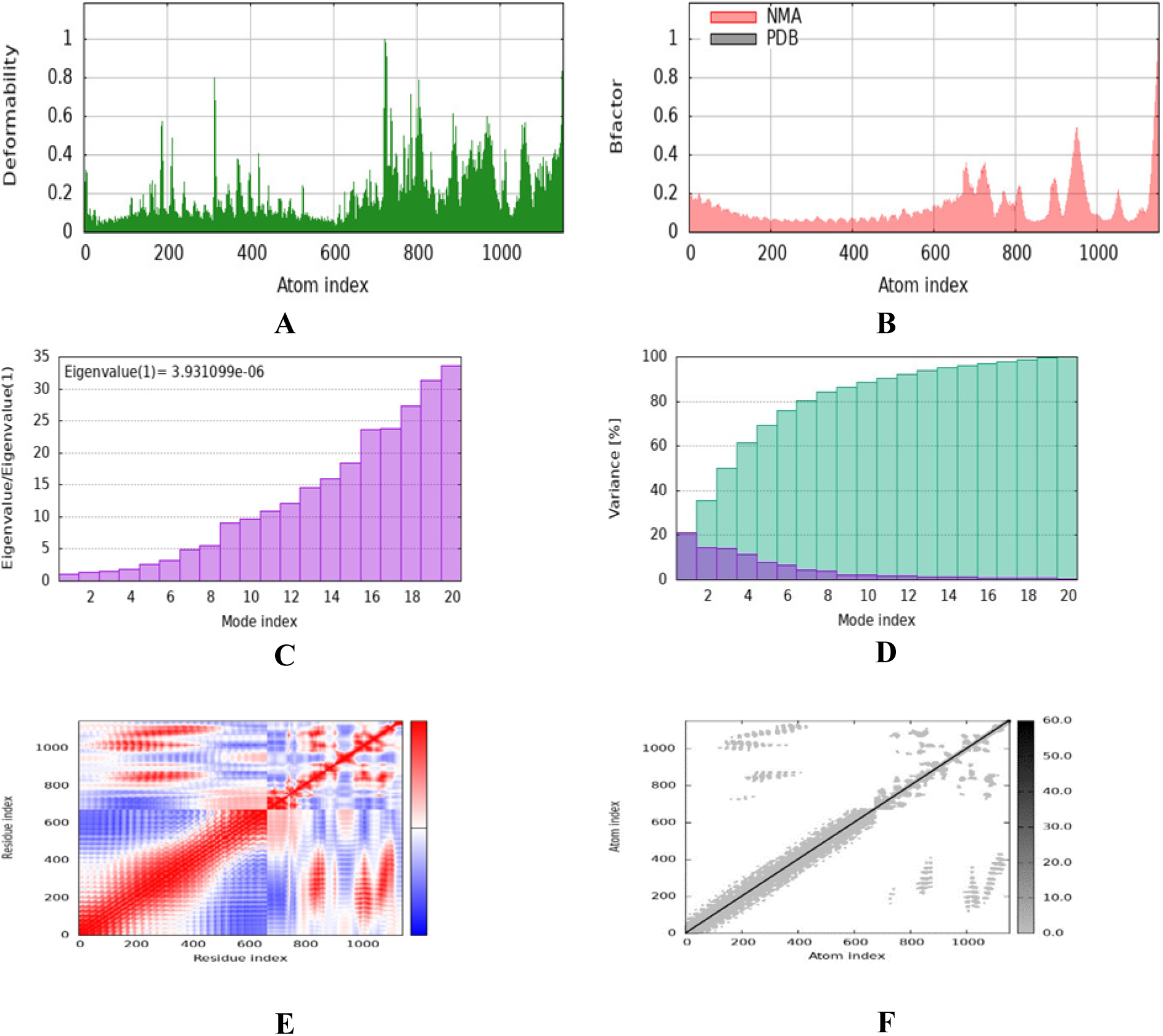
MD Simulations with iMODS server: (**A**) Deformability plot; (**B**) B factor plot; (**C**) variance plot; (**D**) The eigenvalue (**E**) The covariance matrix between pairs of residues where red, white, and blue represent correlated, uncorrelated, anti-correlated motion respectively; (**F**) The elastic network model, with darker gray suggesting more rigid springs.

**Figure 17.**
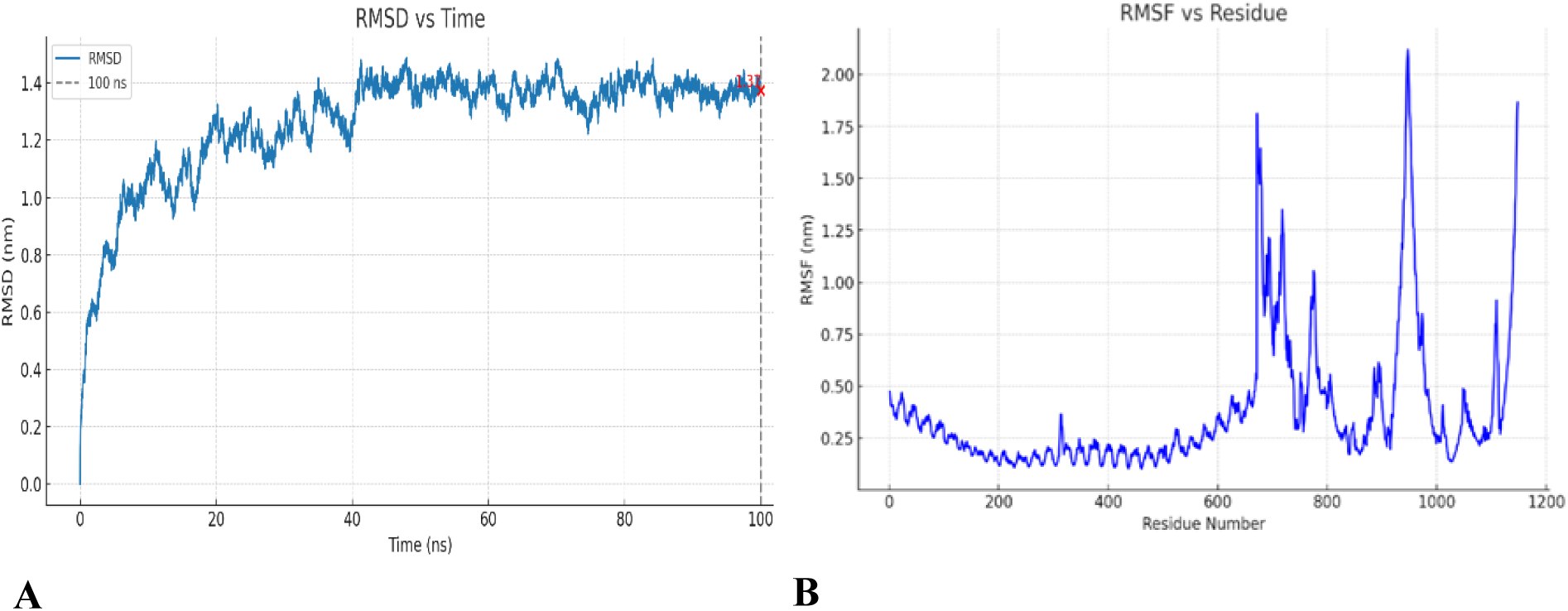
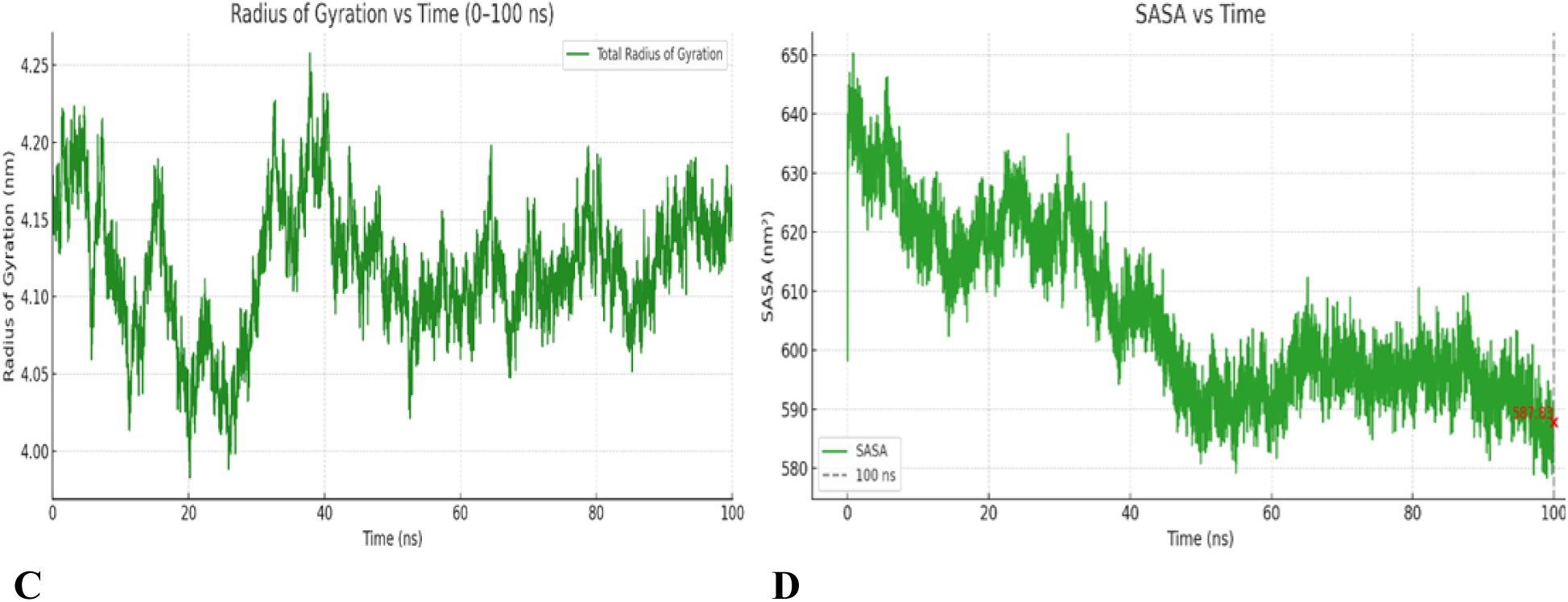
MD simulation of Vaccine contract TLR complex: (A) Root mean square deviations (RMSD) plot, (B) RMSF plot, (C) Rg plotted against simulation time D) The SASA values of the complex was analysed to evaluate changes in the surface of complex.

**Figure 18.**
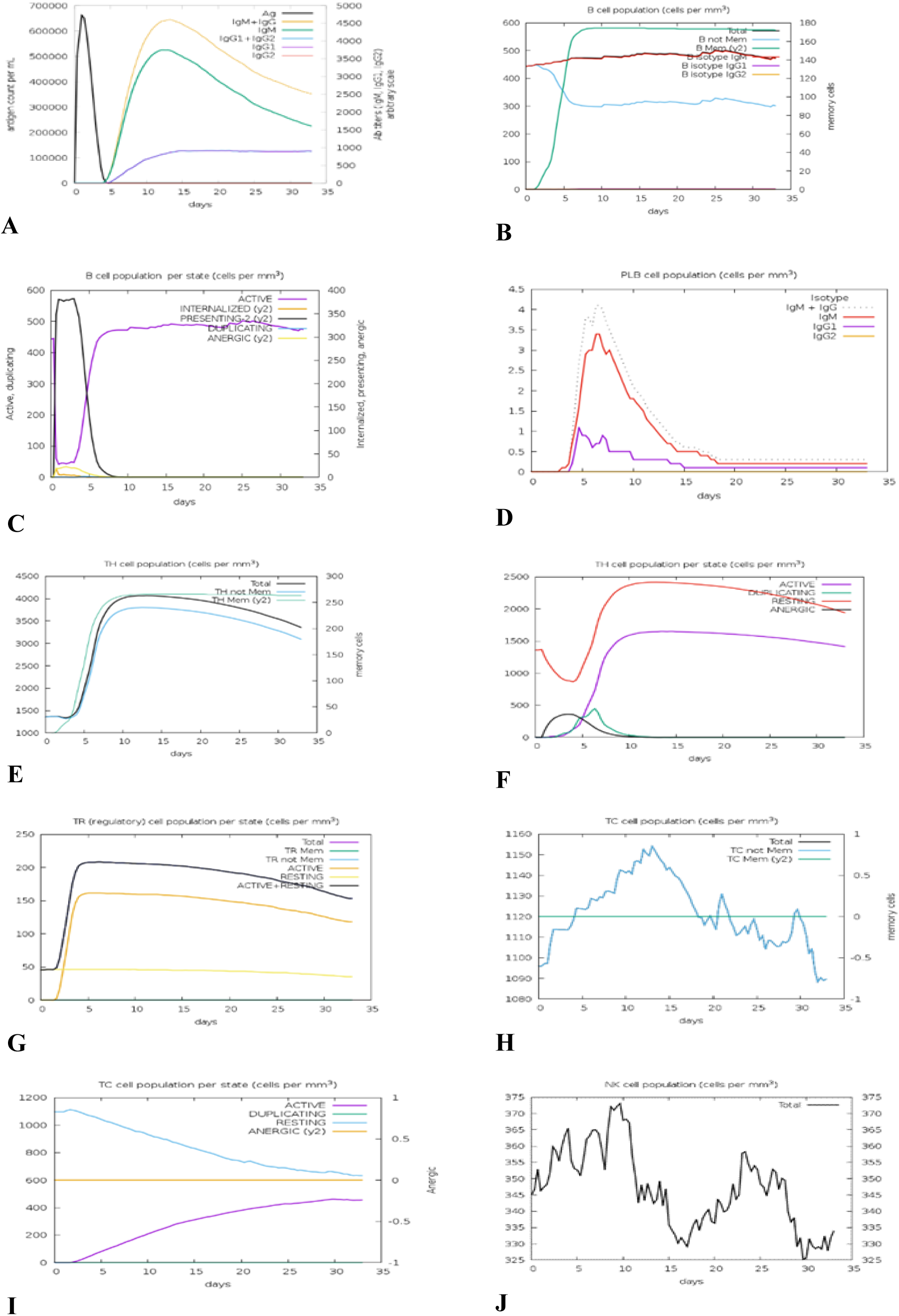
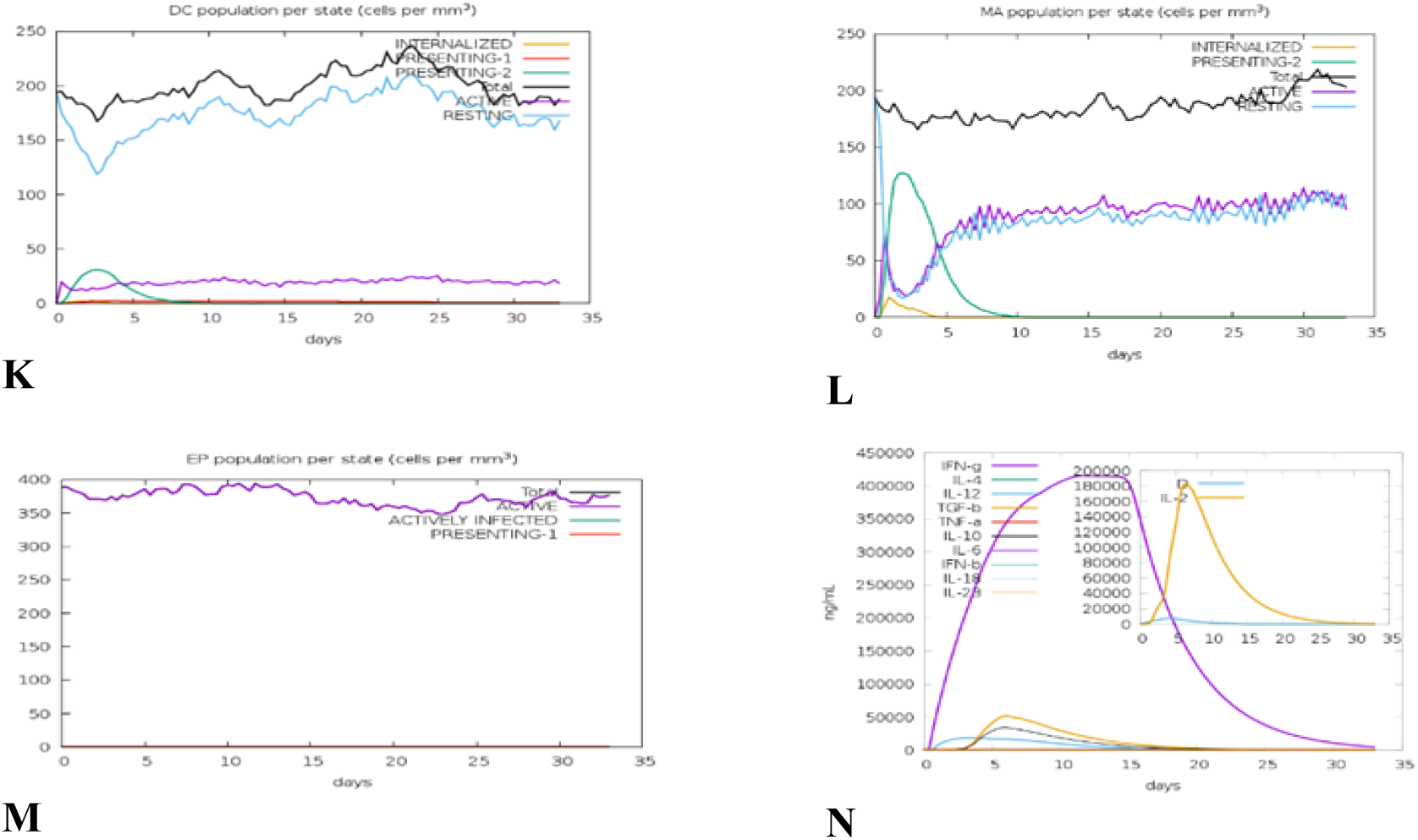
(A) Antigen interaction with immunoglobulins; (B) Activation of B lymphocytes; (C) Proliferation phase of B lymphocytes; (D) Differentiation into plasma B cells; (E) Quantification of CD4+ T-helper cells; (F) Activation of CD4+ T-helper cells; (G) CD4+ regulatory T-cell population; (H) Enumeration of CD8+ cytotoxic T cells; (I) Functional activity of CD8+ cytotoxic T lymphocytes.

**Figure 19.**
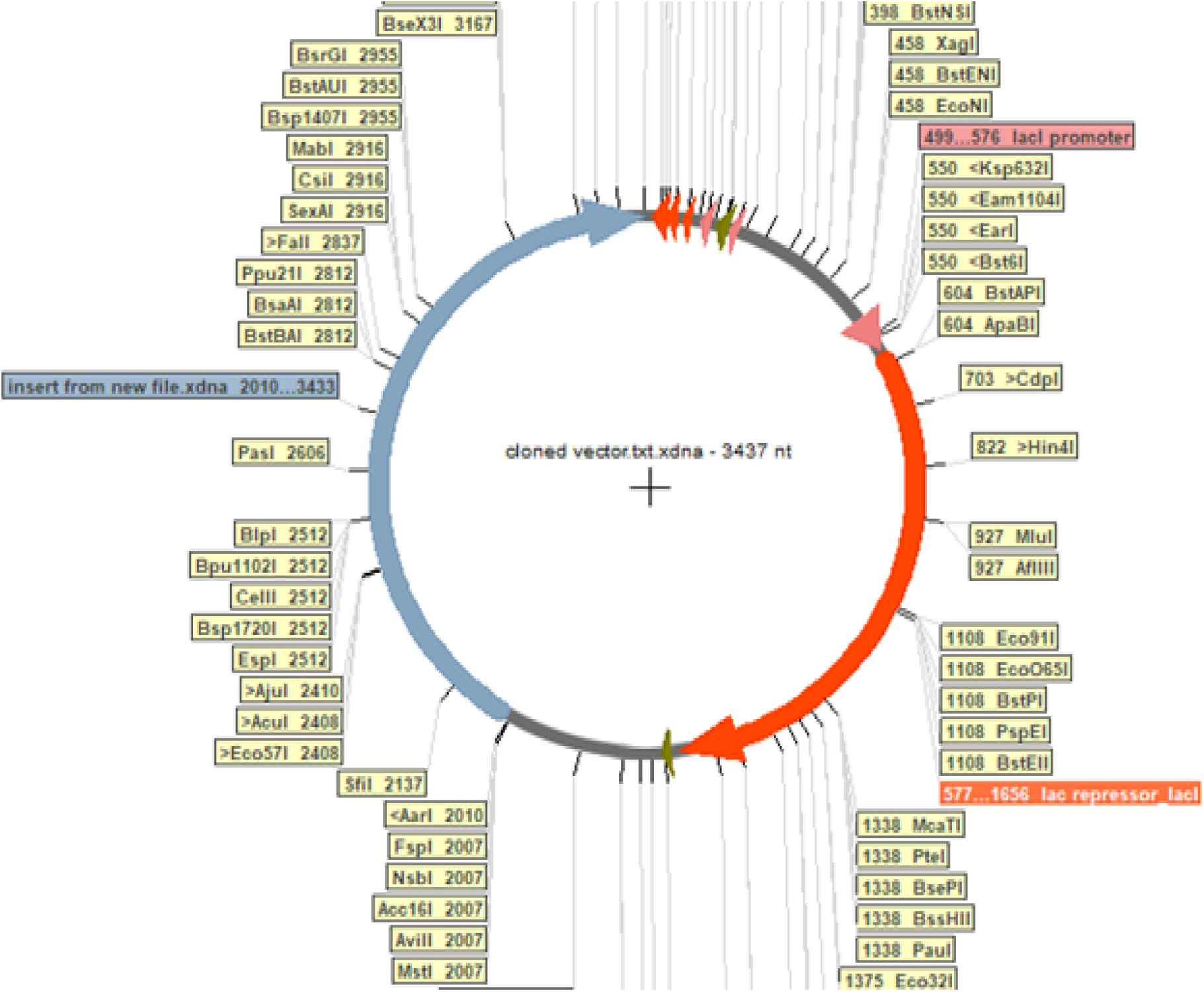
Illustration of in silico insertion of the multi-epitope vaccine construct into the pET28a(+) expression vector using EcoRI and BamHI restriction sites.

### Conformational Stability and Flexibility

MD simulations were carried out on the top-ranked complexes with high affinity for binding (Table 5). The root mean square deviation (RMSD) tracked conformational changes from the initial structure throughout the simulation. The RMSD values started low and increased before stabilizing at 1.37 nm at 100 ns, indicating a significant but stable conformational shift from the starting structure (Figure. 14a).

The RMSF analysis of the complex showed residue fluctuations mostly within 0.5 nm, with increased flexibility observed at interaction interfaces between the proteins (Figure. 14b). The Rg, which reflects protein compactness and folding dynamics, remained stable around 3.65 nm with minor fluctuations of ±0.09 nm throughout the 100 ns simulation, indicating maintenance of structural compactness and equilibrium (Figure. 14c).

The Solvent-accessible surface area (SASA) analysis revealed initial higher values that stabilized over time, fluctuating around 587.83 nm² at 100 ns, reflecting a consistent solvent exposure and further supporting the structural stability of the complex during the simulation (Figure. 14d).

### Predicted Disulfide-bonds

The vaccine construct’s amino acid sequence was examined for disulfide bonds. A Support Vector Machine-based classification method predicted disulfide bonds in the sequence. Out of 5 cysteine residues sequence, 4 (Cys111, Cys130, Cys158, and Cys193) are paired to form 2 disulfide bridges, which are visualized (Figure X) as connecting loops. These covalent linkages likely stabilize the protein’s folded structure by anchoring specific regions together.

*Amino Acids:*

DPNAPKRPPSAFFLFCSEEAAAKWQNNNQTNTNYSGFAPNTLAKHPRYRGTNDTRNPWVNDSLAPDWDNMTWQEWGFAPTKEKRYSSAHGRHFAAPGKGSDPEVARATRECPGPGAMGAASLTATIPLFCATRFVLGFLGFLIIVAVIALRILLASACLVYIQYGVLIIVNMTWFLNWIRLRKGYRPVRQAWCWFKGKRSFLTLQLIYVTSIIANIDWYIQYGVLIIVAAYAASLTVSAQSRTLLAGFLATAGSAMGAASLGFLGFLATAGSAMGALGFLATAGSAMGAASLGFLGFLATAGSAMGLLSRSFLTLQLIYQNLTSWVKYIQYGVLIINQTNITFSAEVAELYPIAYIHFLIRQLIRLRSFLTLQLIYQNLRDSRSFLTLQLIYQNLRSWVKYIQYGVLIIVATSWVKYIQYGVLIIVWIQEAFQAAARATREWPWPIAYIHFLIRQLYGVLIIVAVIALRIVHHHHHH

### Immune Simulations response

C-IMMSIM was used with default parameters except host HLA selection. From the IEDB analysis resource, HLA-A0101, HLA-A0301, HLA-B1501, HLA-B0801, HLA-DRB11101, and HLA-DRB10101 were selected for their high genotypic frequencies in the global population. The immune simulation showed strong balanced in humoral/cellular immune responses. Rapid B cell proliferation, memory cell formation, and plasma cell differentiation with elevated IgM, IgG1, and IgG2 antibodies also witnessed. Both CD4⁺ T-helper and CD8⁺ CTLs showed increased memory and total populations, activation, proliferation. Antigen presentation and immune regulation were also performed by regulatory T cells, natural killer cells, macrophages, dendritic cells, and epithelial cells. A strong immune response was confirmed by high cytokine and interleukin levels, including danger signals.

### Codon Optimization, Expression Vector Selection, and *InSilico* Cloning

Codon optimization was carried out as part of the in silico vaccination development process to improve expression of the vaccination construct in a prokaryotic host. With a Codon Adaptation Index (CAI) of 0.628 and a GC content of 65.48%, the ideal sequence shown good adaptation for expression in Escherichia coli K12. Strong T7 promoter and His-tag sequence drove the pET28(a) vector choice for downstream expression and purification. To allow directional cloning, the gene termini included EcoRI restriction sites and BamHI. Using Serial Cloner, silico cloning confirmed the proper insertion and orientation of the vaccine construct into the pET28(+) vector, so guaranteeing feasibility of experimental expression in bacterial systems.

## Discussion

Although HIV-2 is less prevalent than HIV-1 but is gaining global attention due to its gradual spread beyond West Africa into Asia, Europe, and the Americas, as highlighted by Campbell-Yesufu and Gandhi (2011). In response to the growing need for targeted prevention, this study designed a multi-epitope subunit vaccine aimed specifically at the HIV-2 envelope (env) protein, intending to stimulate both cellular and humoral immune responses.

Multi-epitope vaccines combining CTL, helper T lymphocyte (HTL), and B-cell epitopes, along with potent adjuvants, show strong potential to induce broad cellular and humoral immunity, supported by computational and early experimental data (Rahmani et al., 2019). Our construct incorporated cytotoxic CTL, HTL, and B-cell epitopes selected based on strong immunogenic profiles. Notably, CTL epitopes spanning amino acid positions 30–65 (e.g., 46–55, 30–39, 56– 65) demonstrated high binding affinity. Among HTL epitopes, one located at position 41–55 showed high promiscuity, binding with 11 different HLA class II alleles and yielding a NetMHCpan average rank of 0.716, suggesting robust potential for eliciting both Th1 and Th2 responses.

Population based Global coverage showed promising results for selected MHC-I and MHC-II epitopes covered up to 82% and 99% of the global population, respectively. However, MHC-II coverage in Pakistan was found to be extremely low—just 1%. This likely reflects an underrepresentation of local HLA alleles in global immunoinformatics databases, as noted by Saha et al. (2022), highlighting the need for more region-specific datasets in future vaccine development.

The final vaccine design (477 amino acids; molecular weight of approximately 54 kDa) indicated favorable solubility and high stability. Ramachandran plot analysis confirmed the structural integrity of the model, with 96% of residues located in the most favored regions, indicating a well-folded and stable protein structure are consistent with the results of Meza et al., (2017). Disulfide bond predictions confirmed three stable bridges among five cysteine residues, further enhancing the vaccine’s rigidity and thermal resilience (Wiedemann et al., 2020).

Antigenicity prediction classified the vaccine construct as a strong candidate, with a VaxiJen score of 0.7146—significantly above the accepted threshold of 0.45, indicating robust immunogenic potential. Further computational screening confirmed that the construct is not allergic/toxic, which are essential feature of an ideal vaccine. Immunoglobulin-specific antigenicity scores—IgG (1.217), IgA (0.164), and a negative IgE score (–0.733)—reinforce its likely compatibility with the immune system, especially in minimizing allergic responses. According to Arya et al. (2021), an ideal multi-epitope vaccine should demonstrate high antigenicity, be free from toxicity or allergenicity, and show good solubility and structural stability. These characteristics collectively suggest that Vaccine construct is an excellent candidate.

Among the predicted epitopes, several showed potentials to induce key cytokines: LLSRSFLTLQLIYQN for IL-10 (anti-inflammatory), PIAYIHFLIRQLIRL for IL-4 (Th2-type), and RSFLTLQLIYQNLRD and AMGAASLTV for IFN-γ (Th1-type). One B-cell epitope, GFAPTKEKRYSSAHGRH, was identified as a strong IgG inducer, supporting its role in humoral immunity. Despite these promising features, the low sequence conservation (7.69%) across related proteins suggests possible strain-specific immune responses, which may limit broad vaccine applicability.these results emphasize the potential of the selected epitopes to engage diverse immune pathways while acknowledging the challenges of epitope variability in vaccine development (Nagpal et al., 2017; Dhanda et al., 2013; Bui et al., 2007; Fleri et al., 2017).

Molecular docking analysis discovered favorable interplays between the vaccine and TLRs. The binding energies observed for TLR2 (–1481.1 kcal/mol), TLR3 (–1572.2 kcal/mol), TLR4 (– 1368.5 kcal/mol), and TLR9 (–1669.3 kcal/mol) suggested stable complex formation. Among these, the TLR2 interaction—particularly at the vaccine’s N-terminal—was the most stable, consistent with prior reports that highlight TLR2’s critical role in initiating innate immune responses (Thibault et al., 2009; Browne, 2020).

Our construct also demonstrated that vaccine is capable of evoking strong immunological stimulations. The early rise in IgM, IgG1, and IgG2—starting as early as day 2 and peaking between days 7 and 15—reflects an effective primary response. Notably, repeated exposures led to elevated levels of IgM_IgG_IgG1_IgG2 with cytokines (IFN-γ & IL-2) indicate a potent memory response. The consistently low IL-10 response is indicative of a Th1-skewed profile, which is particularly favorable for antiviral immunity. These findings align with previous immunoinformatics-based vaccine studies, where a Th1-biased immune profile was linked to better control of viral infections (Sher et al., 2023).

Further validation through MD simulations confirmed the vaccine-TLR2 complex’s stability, with RMSD remaining steady around 1.37 nm over 100 ns and low RMSF values indicating minimal fluctuations. A low score of eigenvalues from NMA further explains a rigid, stable interaction. These results align with Abu Tayab Moin et al. (2023), who provide strong binding affinity and consistent hydrogen bonding in TLR-vaccine complexes in his research paper.

The construction’s CAI was nearly 1.0, and GC content within the ideal range for E. coli.The optimized sequence was cloned into the pET28a(+) vector via Serial Cloner 2.1, goals for protein production (Abdulla et al., 2019).

To better perform translational potential, codon optimization was carried out in E. coli. The high CAI (close to 1.0) and optimal GC content illustrate strong potential for efficient expression. The optimized sequence was then successfully inserted into the pET28a(+) vector using Serial Cloner 2.1, for effective protein synthesis (Abdulla et al., 2019).

Compared to earlier HIV-1 vaccine models like the 315-amino-acid construct by Pandey et al. (2018) or the gp120-based epitopes used by Habib et al. (2024) the our design not only longer but also includes a broader range of epitopes and this construct is specifically for to HIV-2, a pathogen that has remained under-estimated in vaccine research instead of its epidemiological importance (Gottlieb, 2013). The enhanced TLR binding profiles, antigenicity scores, and immune simulation outcomes showed significant improvements over previous models. Still, the computational findings are best, they must be done in wet-lab validation. Additionally, the limited MHC-II coverage in populations such as those in Pakistan underscores a pressing need to incorporate local HLA allele data into global immune-informatics pipelines to ensure broader vaccine inclusivity.

## Conclusion

Immuno-informatics-driven design of a multi-epitope vaccine targeting the HIV-2 env protein demonstrates a promising strategy. The construction showed strong antigenicity, structural stability, and broad population coverage while being non-toxic and non-allergenic. In silico analyses streamlined the vaccine development process, reducing cost and time. Although the results are encouraging, experimental validation is essential to confirm the vaccine’s efficacy and immunogenicity in a Lab setting enviremnent.

## Conflict of Interest

The authors declare no conflict of interest.

## Funding

This research received **no specific grant** from any funding agency in the public, commercial, or not-for-profit sectors.

## Supplementary Material

Supplementary material is available upon request.

## Notes

### Competing Interest Statement

The authors have declared no competing interest.

